# Cargo crowding, stationary clusters and dynamical reservoirs in axonal transport

**DOI:** 10.1101/2021.03.12.434740

**Authors:** Vinod Kumar, Amruta Vasudevan, Keertana Venkatesh, Reshma Maiya, Parul Sood, Kausalya Murthy, Sandhya P. Koushika, Gautam I. Menon

**Author notes:** These authors contributed equally to this work (Vinod Kumar, Amruta Vasudevan, Keertana Venkatesh, Reshma Maiya).

## Abstract

Molecular motors drive the directed transport of presynaptic vesicles along the narrow axons of nerve cells. Stationary clusters of such vesicles are a prominent feature of axonal transport, but little is known about their physiological and functional relevance. Here, we develop a simulation model describing key features of axonal cargo transport with a view to addressing this question, benchmarking the model against our experiments in the touch neurons of *C. elegans*. Our simulations provide for multiple microtubule tracks and varied cargo motion states while also incorporating cargo-cargo interactions. Our model also incorporates obstacles to vesicle transport in the form of microtubule ends, stalled vesicles, and stationary mitochondria. We devise computational methodologies to simulate both axonal bleaching and axotomy, showing that our results reproduce the properties of both moving as well as stationary cargo in vivo. Increasing vesicle numbers leads to larger and more long-lived stationary clusters of vesicular cargo. Vesicle clusters are dynamically stable, explaining why they are ubiquitously seen. Modulating the rates of cargo motion-state switching allows cluster lifetimes and flux to be tuned both in simulations and experiments. We demonstrate, both in simulations and in an experimental system, that suppressing reversals leads to larger stationary vesicle clusters being formed while also reducing flux. Our simulation results support the view that the physiological significance of clusters is located in their role as dynamic reservoirs of cargo vesicles, capable of being released or sequestered on demand.

## Introduction

Intracellular cargo, such as membrane-bound organelles and macromolecular complexes, are transported in a directed manner along neuronal processes. Such transport, driven by molecular motor proteins, occurs largely along the bundle of axonal microtubules. Individual motor proteins move in a unidirectional manner, dependent on the intrinsic polarity of microtubules. However, membrane-bound organelles such as mitochondria, endosomes, lysosomes, and synaptic vesicles exhibit bidirectional transport when viewed over long timescales, indicating that both anterograde and retrograde motors participate in transporting neuronal cargo [1]. While endosomes [2] and mitochondria [3] exhibit diverse motility characteristics based on their maturation state and size respectively, much less is known about diversity in synaptic vesicle motility *in vivo*.

Several studies have found that stationary clusters of neuronal cargo are a prominent feature of axonal transport [4–8]. These stationary vesicle clusters have been shown to modulate cargo transport; moving cargo change their state of motion at these locations [8] or slow down in their vicinity [4]. Stationary clusters of synaptic vesicles appear to be associated with actin-rich regions, as well as with microtubule ends *in vivo* [4, 9]. These stationary synaptic vesicle clusters can be dynamic, mobilizing in response to neuronal activity [8]. However, the role of stationary vesicle clusters in maintaining cargo transport along neuronal processes has not been explored in detail, and it is unclear how the neuron modulates the relative fractions of stationary and moving cargo within the axon.

Prior simulation and experimental work has left the question of the statics and dynamics of vesicle clusters unaddressed. Work by Lai, Brown and Xue [10] uses a simple lattice model to suggest that local axonal cargo accumulation can be induced by a global reduction of functional molecular motors in the axon. Yogev and collaborators [11] study the single vesicle version of this model, studying experimentally observed vesicle pausing at microtubule ends in *C. elegans* motor neurons. Rank and Frey [12] study the effects of interactions between Kinesin-1 motors along the microtubules on transport properties and crowding using Monte Carlo simulations, with inputs from experiments.

Apart from these, a number of modelling studies have focused on understanding traffic jams in pathological conditions, for instance, at regions of microtubule polar mismatch [13, 14], and at regions of reduced microtubule densities [15]. Lattice models with multiple lanes and exclusion processes have been used to study transport properties of cargo or motors on microtubule lanes [16–18]. A number of models for motor-driven microtubule-based transport focus on understanding general features of axonal transport as opposed to studying specific model systems [19–24]. However, to the best of our knowledge, no prior work has attempted to account for the diversity of obstacles that moving precursors of synaptic vesicles (pre-SVs) in dense axons might encounter nor the physiological implications of the resultant crowding of moving pre-SVs. Indeed, how is it that cargo can be transported steadily in healthy neurons despite the inevitable encounters of oppositely directed pre-SVs in the dense axonal environment?

This paper combines experimental observations of *in vivo* pre-synaptic vesicle transport in the touch receptor neurons (TRNs) of *C. elegans*, with kinetic Monte Carlo simulations of a model constructed to describe these observations. Our experiments show that moving synaptic vesicles exhibit three distinct motion states, i) smooth anterograde motion, with longer run lengths and few or no pauses, ii) step anterograde motion, with shorter run lengths and more frequent and longer pauses, and iii) retrograde motion. The model accounts for multiple microtubule tracks and prescribes hopping rates for vesicles of each of the three motion states outlined above, consistent with experimentally-derived proportions.

Our simulations further account for cargo crowding, wherein vesicle motion can be blocked in three distinct ways: (i) crowding due to proximity of vesicles on the same track, (ii) the presence of open ends of microtubules, preventing vesicles from translocating further along the same track, and (iii) the presence of large blocks to motion, such as stalled mitochondria, which use the same tracks to translocate. We describe two novel simulation methodologies for the modeling of fluorescence recovery after photobleaching and the accumulation of vesicles following axotomy, developed to compare vesicle transport between experiments and simulations. We then use this simulation model to examine the effect of altered synaptic vesicle motility characteristics on net vesicle transport. Implementing these specific perturbations is straightforward in simulations, although experimentally difficult. We find that relatively small changes in rates of motion state conversion have a large effect on cargo flux and the sizes and lifetimes of stationary vesicles along the axon.

Our simulation and *in vivo* data highlight the importance of microtubule track switching (sidestepping) and bidirectional motion of moving cargo for efficient cargo transport along the neuronal process. This indicates that the ability to tune vesicle motility characteristics, perhaps by activating or inactivating a class of motors consequent to a signaling event *in vivo*, could potentially enhance or suppress net cargo transport [25–27]. We connect these observations to the idea of “dynamic reservoirs” of vesicles at locations of stationary clusters, noting that currents remain steady even as vesicles can be trapped at stationary clusters over a broad range of times. We suggest that the physiological significance of stationary clusters lies in the fact that the net sizes of these clusters can be tuned through small changes in rates at which vesicle states interconvert, providing a fast way to mobilize vesicles from nearby locations upon a signaling event.

## Results

### Characterizing vesicular cargo motion along the *C. elegans* touch receptor neurons provides input to simulations

We characterized the transport of synaptic vesicles along the process of the PLM touch receptor neuron in *C. elegans*. The PLM neuronal process is crowded with microtubules [28], and also likely contain several actin-rich regions [8]. We tracked fluorescently labeled precursors of synaptic vesicles (pre-SVs) using GFP-tagged to RAB-3 [29, 30], finding that moving vesicles exhibited diverse motility characteristics (Fig. S1 A). While synaptic vesicles show both anterograde and retrograde transport, the anterograde-directed population of vesicles showed two distinct motion states, distinguished on the basis of the number of pauses exhibited in the trajectories of individual vesicles (Fig. S1 B). We categorized anterograde-directed vesicles exhibiting fewer than 0.15 pauses/sec as ‘Smooth anterograde (SmA)’ and vesicles showing more than 0.15 pauses/sec as ‘Step anterograde (StA)’. Further analysis revealed that smooth anterograde (SmA) motion was processive, with an average run length of 1.3 ± 0.8 *µm*, and a maximum run length of 8 *µm*, interspersed by brief pauses (0.2 ± 1.8 sec) (Fig. S1 C and D). Step anterograde (StA) motion, on the other hand, is characterized by higher pause numbers, longer pause durations (2.7 ± 3.6 sec), and shorter run lengths (0.8 ± 0.5 *µm*) as compared to SmA (Fig. S1 C and D). We also observe smooth retrograde (R) motion towards the cell body, associated with an average run length of 1.9 ± 1.7 *µm*, and a maximum run length of 14 *µm*, as well as fewer and shorter (0.2 ± 1.5 sec) pauses (Fig. S1 C-D). We only rarely observe step retrograde motion (< 1%) and neglect this possibility to a first approximation. Our experiments show that vesicles exhibit these three motion states in a well-defined ratio of approximately 50 (SmA):10 (StA):40(R) (Fig. S1 E).

In addition to the anterograde and retrograde motion states, we also observe that pre-SVs can change their direction of motion along their trajectory. We term this a ‘reversal’ (Fig. S1 A). We observe that an average of 13.68% (1.368 ± 0.47717 per 10 RAB-3 vesicles) of total moving RAB-3 vesicles reverse along the PLM process and that both anterogradely and retrogradely moving vesicles show reversals along their trajectory (Fig. S1 G). Further, a population of pre-SVs are stationary for a broad distribution of timescales along the neuronal process, with several being stalled for over 2-3 minutes (Fig. S1 F). Based on the time duration for which vesicles remain stationary, we define stationary vesicle clusters as ‘short-lived stationary clusters’ and ‘long-lived stationary clusters’ (described in detail in the ‘Methods’ section). In TRNs, long-lived stationary vesicle clusters of RAB-3 contain 2-4 vesicles [8]. We see no evidence for predominantly diffusive motion of pre-SVs along the PLM neuronal process, which is discussed in subsequent sections. We believe this is because *C. elegans* touch receptor neurons are crowded with microtubules [28]. A microtubule-dense process would promote directed transport due to repeated encounters of the cargo-motor complex with microtubules, compared to the case of a less dense axoplasm. These prominent experimental features are incorporated into our simulation model.

### Simulations reproduce properties of vesicle transport across multiple scales

We compare vesicle trajectories generated in our model, motivated by experiments, to vesicle trajectories in *C. elegans* neurons. We choose hopping and state inter-conversion rates to reproduce both the mean velocities as well as the steady state ratios of StA, SmA and R in experiments. Quantities such as mean run lengths, pause durations and pause frequencies follow from these choices. Fig. 1 A and B provides a snapshot of model-derived vesicle trajectories, with rates described in Table S2. The snapshot in Fig. 1 A shows a configuration of a “simple model” in which there are no blocks to transport, apart from those which form when moving vesicles encounter each other.

**Fig 1.**
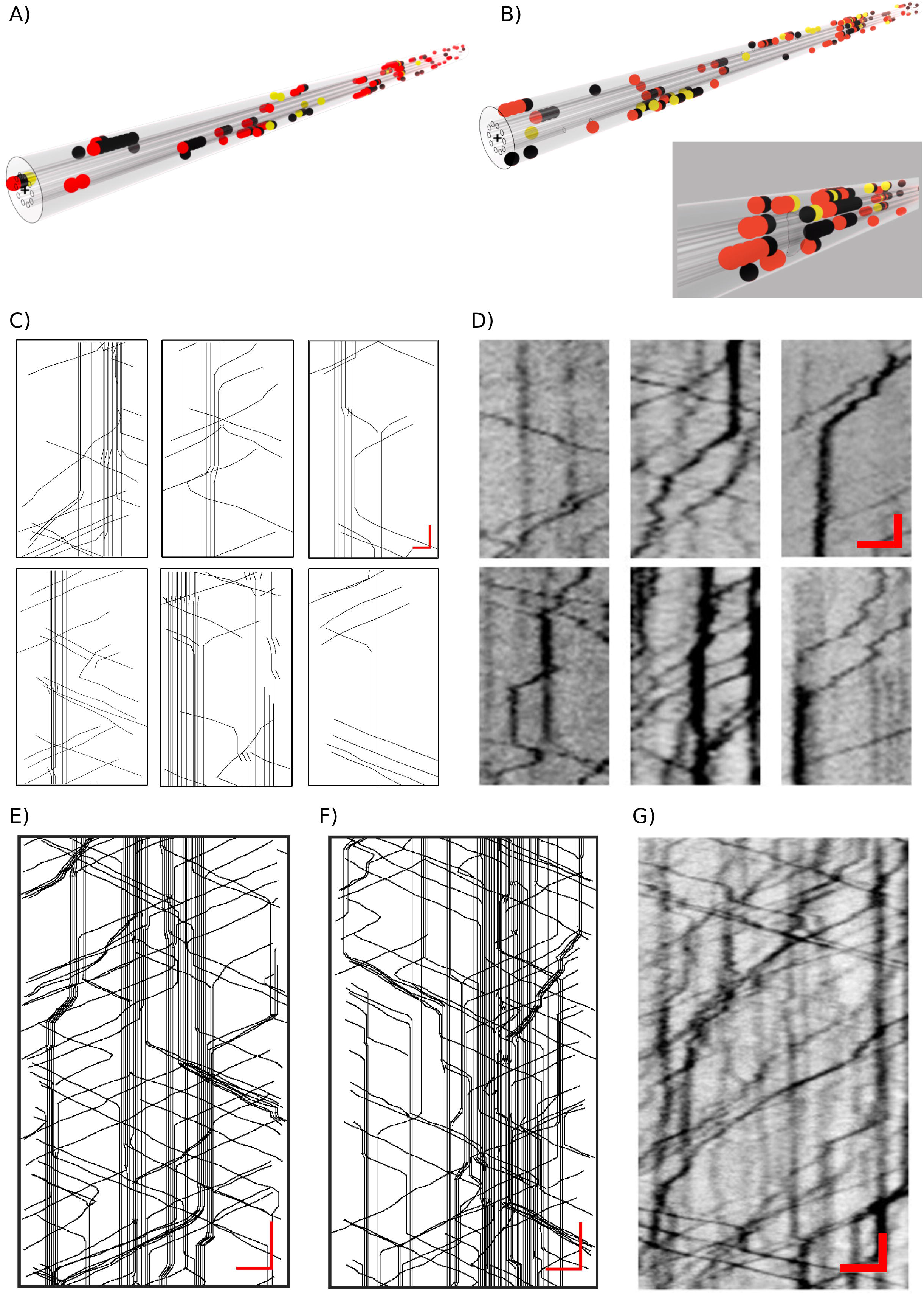
Simulations reproduce key features of cargo transport in vivo. (A) Snapshot of vesicle positions, colour coded as StA (yellow), SmA (black) and R(red), in the case where there are no microtubule breaks or other slow-moving cargo to obstruct motor-induced vesicle motion. The few stationary vesicle clusters that we see arise from the collisions of oppositely directed vesicles, (B) Snapshot of vesicle positions in the case where we allow for a number of extrinsic blocks to vesicle motion, including 6 microtubule breaks leading to 12 ends, and one mitochondrion. Here we see far more stationary vesicle clusters, most of which correlate strongly to microtubule ends. (inset to B) A snapshot of a “stuck” or “blocked” configuration, in which net vesicle motion ceases and there is typically one large stationary cluster. Such states appear to be obtained at vesicles densities that are not too high, provided vesicles are only allowed to change their track and not their state. Examples of trajectories involving vesicle motion in simulations (C) and in vivo experiments (D), including motion across a stationary cluster of vesicles, the unblocking of a two-vesicles stationary cluster (dispersion), cluster formation, the transition between stuck and mobile states of individual vesicles, “crosstalk” between neighbouring clusters of stationary vesicles and the augmentation of a pre-existing large cluster through the addition of one more vesicle. For the simulation kymograph snapshots, the x-axis scale bar = 0.1 *µm* and the y-axis scale bar = 0.5sec. In the experimental kymograph snapshots, the x-axis scale bar = 3 *µm* and the y-axis scale bar = 3sec. In (E), we show an example of simulation kymographs for the simple system, where clusters of stationary vesicles as well as moving trajectories corresponding to the three main vesicles types we consider are displayed. Scale bar: x axis = 2 *µm*, y axis = 1sec. In (F), we show a representative simulation kymograph for the complex system. Scale bar: x axis = 2 *µm*, y axis = 1 sec. For comparison, in (G) we show a sample kymograph from experimental data. Scale bar: x-axis = 3 *µm*, y-axis = 3sec.

The snapshot in Fig. 1 B, is from a simulation of a “complex model” in which we incorporate obstacles to transport. These obstacles are in the form of 6 breaks in the microtubules, corresponding to 12 ends and one mitochondrion spanning 3 microtubules, over 1000 sites along the microtubule lattice. These numbers are comparable to known densities of microtubule ends and stationary mitochondria in the TRNs, and serve to illustrate those aspects of vesicle crowding at obstacles that are relevant to this study [27, 28, 31, 32]. The inset below Fig. 1 B supplies an expanded view of the region proximate to the mitochondrion, showing a number of vesicles stalled in its proximity.

We generate kymographs for the simple system and the complex system, showing representatives of these in Fig. 1 C-F. These kymographs are calibrated to the data and represent the same real-time information as in experiments. We note that they are visually similar to kymographs generated from imaging synaptic vesicles in *C. elegans* TRNs (Fig. 1 D and G). Several features of *in vivo* cargo transport, such as i) moving vesicles halting to form stationary vesicle clusters, ii) vesicles mobilizing from stationary clusters, and iii) moving vesicles crossing stationary clusters, are reproduced in our models of simple and complex systems (Fig. 1 C and D). Both the simulation and the experimental kymographs exhibit prominent clusters of stationary vesicles. The visual similarity of kymographs generated from simulations and experiments (Fig. 1 E-G), as well as the identification of common microscopic properties of moving vesicles, indicates that our model captures significant features of axonal transport in *C. elegans* TRNs.

### Recovery profiles after photobleaching are similar in simulations and experiments

Fluorescence recovery after photobleaching (FRAP) experiments allow investigations into the nature and rates of transport processes [33]. Recovery occurs when unbleached fluorophores, associated here with vesicular proteins, transit into the bleached region. If the region that is bleached contains stationary clusters, fluorescence recovery is also affected by the dynamics of vesicles that halt while moving across them. Experimentally, we examine the recovery profile of vesicle fluorescence after photobleaching in *C. elegans* TRNs, comparing it to the recovery profiles predicted by our models. Since mapping the absolute scale of fluorescence intensity to numbers of labelled vesicles is not straight-forward, we concentrate on understanding the time-scales of recovery.

Our simulation system represents a 64 *µm* section of the axon with 400, 800 and 1000 vesicles. In our simulations, we bleach an 8 *µm* section of the axonal compartment and track individual vesicles that subsequently move within and across the bleached region. Fig. 2 A shows representative bleach recovery kymographs for densities of 400, 800 and 1000 vesicles respectively, which are visually similar to the experimental bleach recovery kymograph. We find that in the simple system, the extent of fluorescence recovery is 80% at lower vesicle densities (400 vesicles). This decreases to 60% at higher vesicle densities (1000 vesicles) (Fig. 2 B-D). In the complex system, while the extent of recovery is lower than that of the simple system, a similar trend is observed (Fig. 2 E-G). At higher densities, vesicle-vesicle interactions and stationary cluster formation likely impede the extent of recovery. Bleached vesicles can linger in the bleached region if stalled, presenting a barrier to the free motion of the unbleached ones. This explains the reduced recovery in the complex system when compared to the simple system. The presence of stationary vesicle clusters in the vicinity of the bleached region also reduces the entry of vesicles into the bleached region, leading to slower recovery. While the timescales of recovery in both the simple and complex systems parallel those observed in experiments, the complex system with relatively high vesicle density provides results that are closest to the fluorescence recovery profiles observed *in vivo* (Fig. 2 H and I). This suggests that *C. elegans* touch neurons present a crowded, vesicle-dense environment, in which moving cargo encounter numerous obstacles to motion in the form of long-lived stationary vesicle clusters, microtubule ends, and other stalled/slow-moving cargo such as mitochondria.

**Fig 2.**
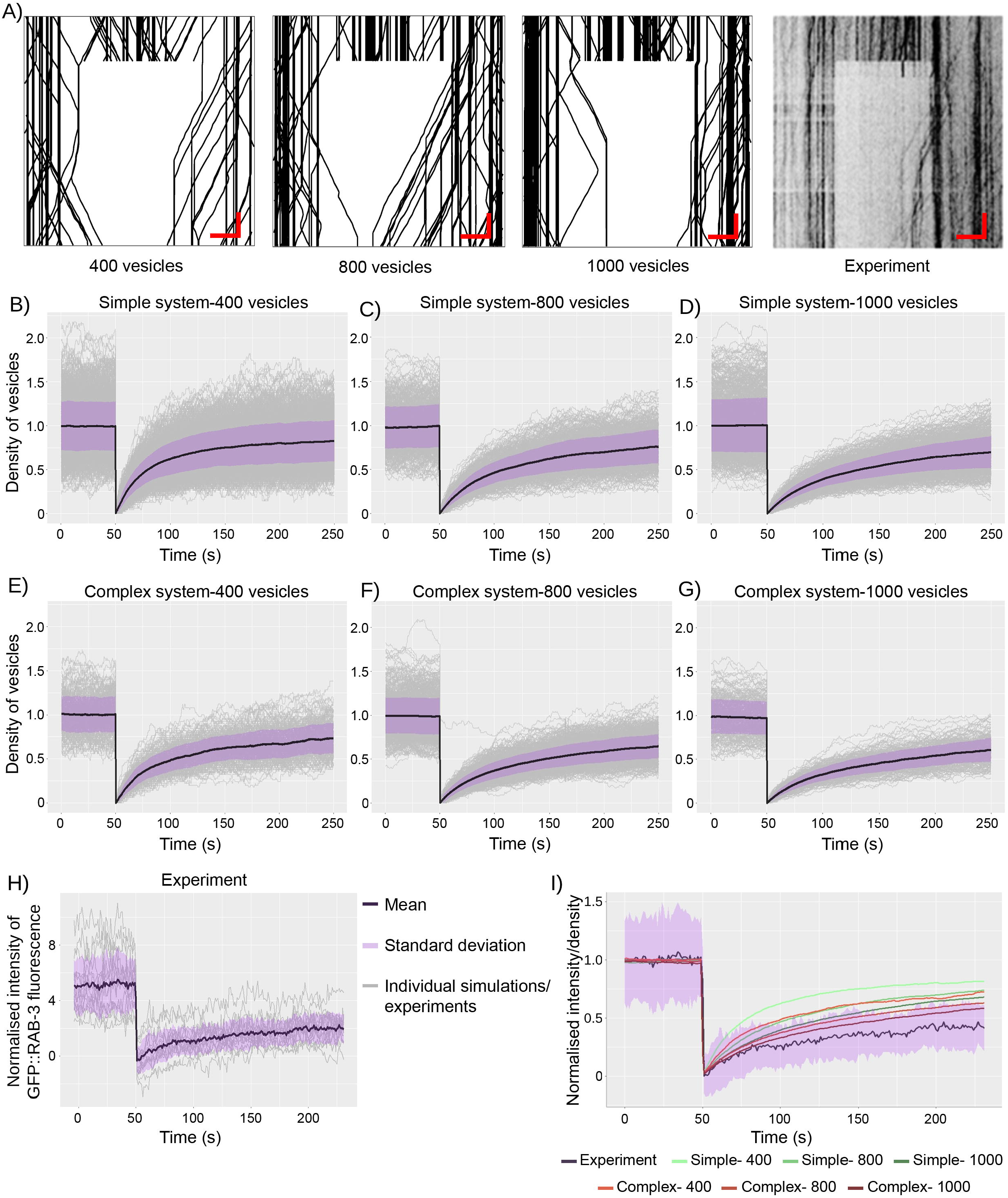
Simulations reproduce the extent of vesicular fluorescence recovery after photobleaching observed in *C. elegans* touch receptor neurons. (A) Representative kymographs illustrating bleach recovery over an 8 *µm* region in a system with microtubule ends and mitochondria, for vesicle densities of 400, 800, and 1000 vesicles, compared to an experimental kymograph illustrating bleach recovery. Scale bar for the simulation kymograph: x-axis = 2 *µm*, y-axis = 1sec. Scale bar for the experimental kymograph: x-axis = 3 *µm*, y-axis = 3sec. (B)-(I) The plots show normalized density of vesicles (along the y-axis) plotted against time (along the x-axis). The fluorescence recovery is monitored for 3 minutes post-bleaching. The grey lines depict fluorescence recovery obtained in individual simulation runs (>100 for each condition), while the black line represents the average of all simulation runs and the purple band represents the standard deviation. (B, C, D) The plots demonstrate the extent of bleach recovery observed in the simple system with an initial vesicle density of 400, 800 and 1000 vesicles respectively. Data is collected from 977, 491 and 723 simulations respectively. (E, F, G) The plots demonstrate the extent of bleach recovery observed in the complex system with an initial vesicle density of 400, 800 and 1000 vesicles respectively. Data is collected from 200, 651 and 228 simulations respectively. (H) The plot demonstrates fluorescence recovery observed in experiments conducted on *C. elegans* touch receptor neurons (N (number of animals) = 16 animals). (I) The plot depicts the average fluorescence recovery profiles for each simulation condition, compared to the experimental fluorescence recovery profile. The standard deviation of the experimental data is depicted as a purple band. Only the fluorescence recovery profile of the complex system with the highest vesicle density (1000 vesicles) falls within one standard deviation around the mean of the experimental data.

### Simulations reproduce accumulation of vesicles following axotomy

Vesicle transport rates along axonal processes can also be probed by invasive methods, including laser-induced ablation of the axon (axotomy). As a consequence of continuous transport, after experimental axotomy, vesicles begin to pile up at the injury sites, with anterogradely moving vesicles enriched at the proximal cut site (closer to the neuronal cell body) and retrogradely moving vesicles enriched at the distal cut site (closer to the neuronal tip). As described in methods, we treat axotomy in our simulations as being equivalent to introducing a permanent block to vesicle motion at two locations across all microtubules, separated by a physical distance of 0.8 microns (100 sites), and removing all vesicles and microtubules that lie within the ablated region.

Representative kymographs of simulated axotomy in the simple system show enrichment of vesicles at both the proximal and distal cut sites (Fig. S2 A-C). The accumulation of vesicles increases with the density of vesicles. It is found to be consistently higher at the proximal cut site, compared to the distal one (Fig. S2 A-C). A similar trend is observed for the complex system, although the extent of vesicle accumulation at the cut sites is lower, as compared to the simple system (Fig. S2 F-H). We monitored the accumulation of vesicles at the proximal and distal cut sites over time and found that, in both the simple and complex systems, the proximal cut site showed a consistent increase in accumulation over time (Fig. 3 A and B, Fig. S2 D and E and Fig. S2 I and J). In contrast, the accumulation at the distal cut site showed a less prominent increase (Fig. 3 A and B). The increase in intensity at the proximal cut site is also augmented by an increase in the number of sites where vesicles accumulate near the site of the axotomized region (Fig. 3 A and B). On moving away from the site of axotomy, the density of vesicles was observed to rapidly decay. This is consistent with vesicle accumulation profiles observed from experiments in *C. elegans* touch neurons (Fig. 3 C and D). We conclude that the comparison between simulated axotomy and the experiments indicate that the model captures key features of the experiments.

**Fig 3.**
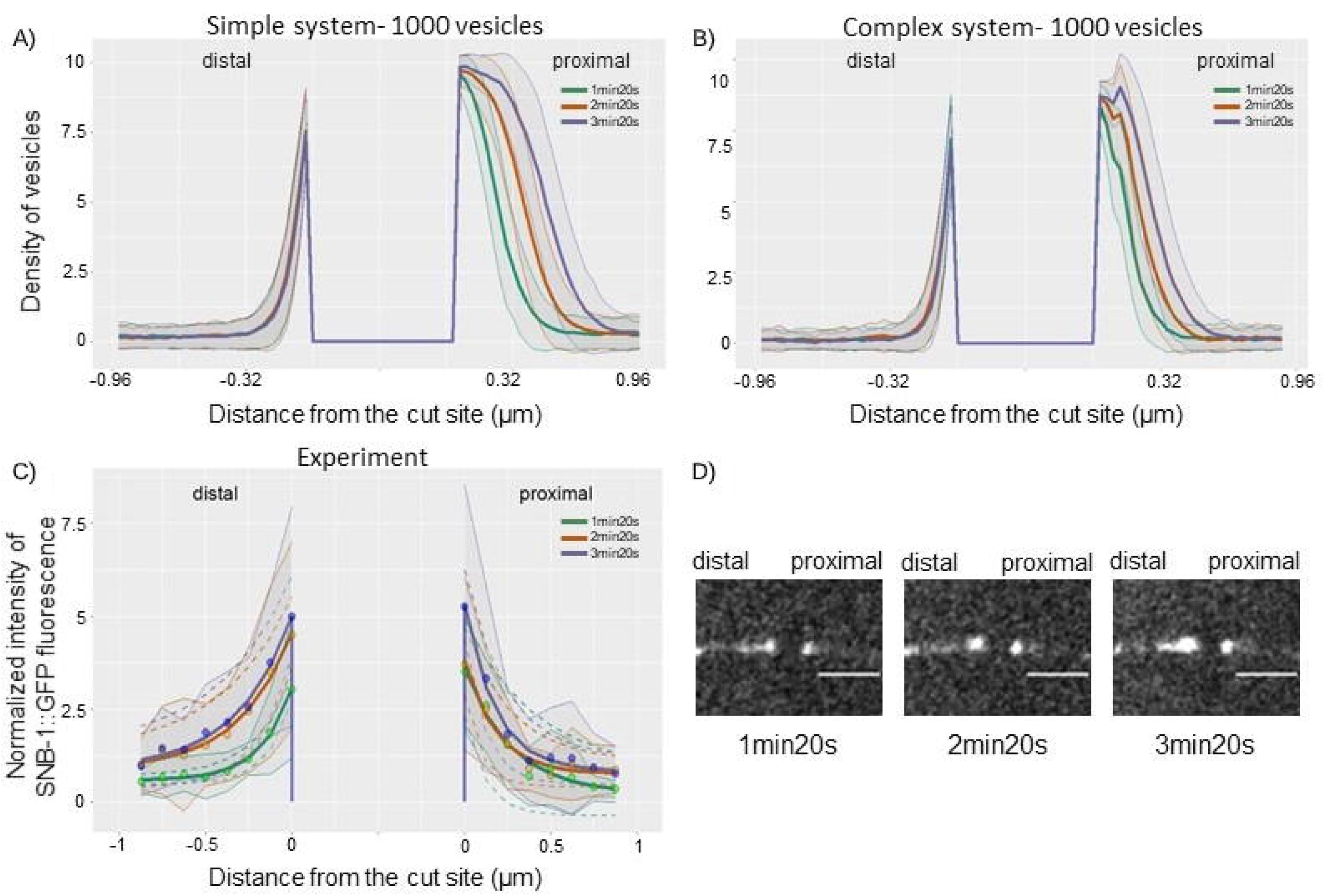
Simulations reproduce vesicle accumulation profiles observed after axotomy *in vivo*. (A) The plot represents the density profile of vesicles 1 *µm* on either side of the site of axotomy for a system of 1000 vesicles moving in a smooth background lacking microtubule ends and mitochondria (simple system), with an axotomized region of 0.8 microns (100 sites in our simulations). The mean density profiles are calculated at 1min20 seconds, 2min20 seconds, and 3min20 seconds post-axotomy, with standard deviation for each plotted as grey bands. Data is collected from 470 simulations. (B) The plot represents the mean density profiles for a system with 6 microtubule ends and 1 mitochondrion (complex system), with an axotomized region of 0.8 microns (100 sites in our simulations) and initial density of 1000 vesicles. Data is collected from 134 simulations. (C) The plot represents mean normalized intensity profiles from axotomy conducted in *jsIs37* transgenic animals (N = 10), with an axotomized region of 3 microns. The mean density profiles are calculated at 1min20 seconds, 2min20 seconds, and 3min20 seconds post-axotomy. (D) The panel shows representative images of the proximal (right) and distal (left) cut sites at (i) 1min20 seconds, (ii) 2min20 seconds, and (iii) 3min20 seconds post-axotomy of the *C. elegans* PLM neuron. Scale bar = 5 *µm*.

### Vesicle density, microtubule ends, and stalled mitochondria regulate the formation of long-lived stationary clusters

We examined the effect of total vesicle numbers and presence of obstacles on the distribution of vesicles along the neuronal process, using our simulation model. Fig. 4 shows time-averaged density profiles for vesicles across 10 microtubules with 1000 sites (8 *µm* region) each, with 50, 100, and 200 vesicles. Obtaining absolute vesicle numbers in experiments is difficult since there is no accurate method to estimate the number of vesicles in stationary cargo. However, we believe that this range should cover the variation in vesicle numbers observed *in vivo*. In panels A, C and E, for the simple system, the densities of vesicles across all sites in SmA, StA and R motion states are small, and present a roughly flat profile. In the simple system, an increase in overall vesicle density from 50 to 100 vesicles increases the baseline vesicle density at all sites along the simulated axon (Fig. 4 A and C). Further, the roughness of the time-averaged density plot provides evidence for the spatial randomness in stationary vesicle cluster formation. More extensive averaging restores the spatial uniformity of density, since such local density peaks have a finite lifetime and can form anywhere.

**Fig 4.**
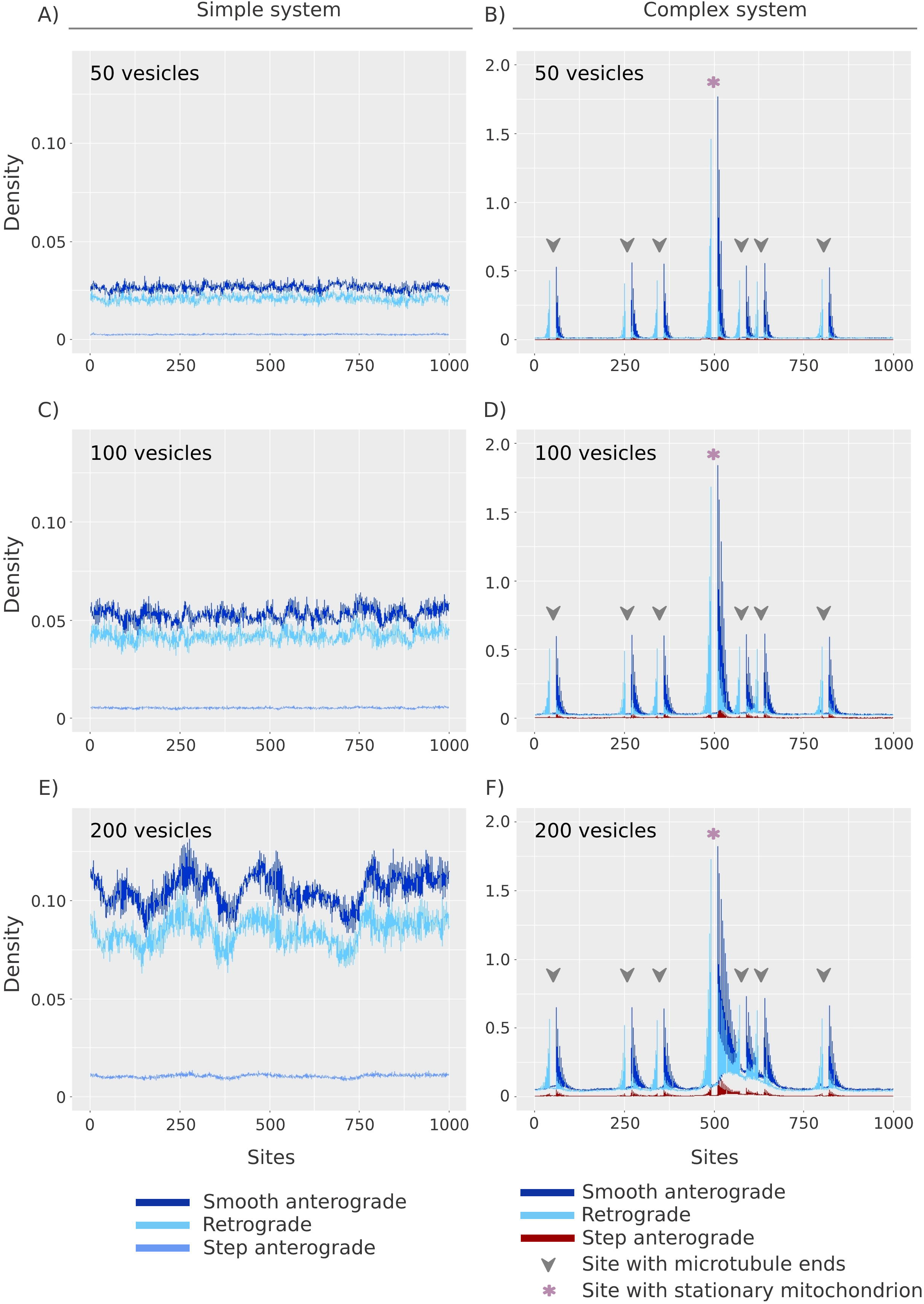
Vesicle density and physical obstacles to transport influence the location and extent of formation of stationary vesicle clusters. (A) Density profiles for SmA, StA, and R vesicles, for a total of 50 vesicles, in a system without microtubule ends or mitochondria (simple system), over 1000 sites. (B) Density profiles for SmA, StA and R vesicles, for 50 vesicles, with 6 microtubule ends and 1 mitochondrion (complex system), with rates as indicated in Table S2. (C) Density profiles for SmA (blue), StA (green) and R (red) vesicles, for 100 vesicles, without microtubule ends or mitochondria. (D) Density profiles for SmA, StA and R vesicles, for 100 vesicles, with 6 microtubule ends and 1 mitochondrion, with rates as indicated in Table S2. (E) Density profiles for SmA, StA and R vesicles, for 200 vesicles, without microtubule ends or mitochondria. (F) Density profiles for SmA, StA and R vesicles, for 200 vesicles, with 6 microtubule ends and 1 mitochondrion, with rates as indicated in Table S2.

Fig. 4 B, D, F describe the effects of introducing obstacles in the form of 6 microtubule ends and a stalled mitochondrion in our model, i.e. the complex system. While both the simple and complex systems reproduce several experimental features of synaptic vesicle transport (Fig. 1 C and D), the complex system is better able to reproduce the distribution of long-lived stationary vesicle clusters observed in experiments (Fig. 1 F, Fig. S3 A). While the simple system shows no spatial bias in the formation of stationary vesicle clusters, as evidenced by the flat density profiles (Fig. 4 A, C, and E), we show that long-lived vesicle clusters, seen as peaks in the density profile, are preferentially formed at locations of obstacles in the complex system (Fig. 4 B, D, and F). Our models suggest that at the lowest vesicle densities, relatively few vesicles stall and form clusters (Fig. S3 A). As the vesicle density is increased, even without the presence of microtubule ends and mitochondria, short-lived clusters are formed (Fig. S3 A and B). With the introduction of microtubule ends and mitochondria, long-lived clusters form at these locations, even if no individual vesicle is permanently bound to those regions. These clusters consist of dense accumulations of vesicles, where vesicle motion is constrained by the fact that vesicles interact, leading to crowding. This phenomenon has also been observed *in vivo* along the neuronal processes of *C. elegans* touch receptor neurons, wherein the presence of stationary vesicle clusters at actin-rich regions increases the propensity of stalling of moving cargo by > 5-fold as compared to actin-rich regions without stationary vesicle clusters [8]. However, long-lived stationary vesicle clusters also form in the simple system, especially at high vesicle densities (Fig. S3 A and B). This suggests that steric interactions arising only from stalled vesicles or vesicles stalled at regions with cytoskeletal heterogeneities are sufficient for the formation of long-lived stationary clusters. This stalling occurs even in the absence of other obstacles to transport, and such clusters of stationary vesicles can themselves serve as roadblocks to transport.

### Small changes in rates lead to large changes in numbers of stalled vesicles and cargo flow

We propose that the conversions between motion states (StA, SmA, R) exhibited by moving vesicles, determine the properties of long-lived stationary cargo clusters. Since it is experimentally infeasible to specifically change the proportion of vesicles exhibiting different motion states, we use our simulations to examine changes in stationary cluster formation and vesicular flux when the rates of interconversion between different motion states are reduced.

To test the effect of altered rates on cargo transport in our simulations, the rate of conversion from Smooth anterograde (SmA) to Step anterograde (StA) was reduced to a third of its earlier value, keeping other rates the same. Fig. 5 A shows representative kymographs from the simple and complex systems for regular and reduced rates of switching between motion states. It can be clearly seen that when rates of switching are reduced, there is a drastic reduction in net cargo flow (Fig. 5 C) and a greater proportion of vesicles are associated with long-lived stationary vesicle clusters (Fig. 5 B). We corroborate this result by quantifying the total proportion of stalled vesicles associated with short-lived and long-lived stationary vesicle clusters (Fig. 5 B, Fig. S3). The figure also shows the percentage distribution of all stationary vesicle clusters across various time windows for initial densities of 50, 100 and 200 vesicles at regular rates and reduced rates of switching.

**Fig 5.**
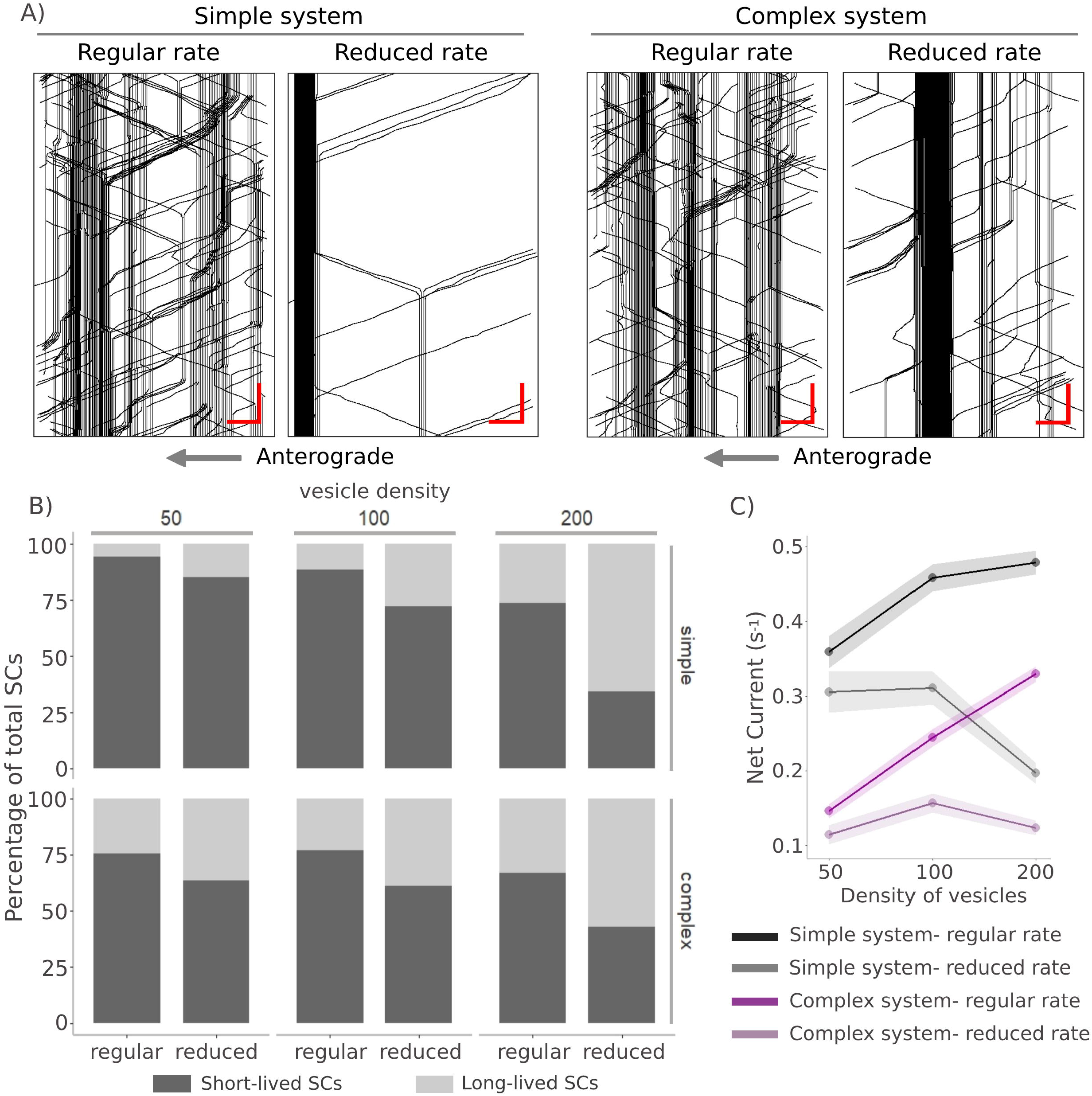
Rates of switching between smooth and staggered motion states influences cargo flow and stationary cluster lifetimes. (A) Representative kymographs for the simple and complex systems with regular and reduced rates of switching between motion states. Scale bar: x axis = 0.5 *µm*, y axis = 2 sec. (B) The panel shows the proportion of long-lived and short-lived stationary vesicle clusters across different vesicle densities at regular and reduced rates for the simple and complex systems. Total number of stationary vesicle clusters are pooled across 150 simulations for each condition. (C) The panel shows net anterograde current across different vesicle densities at regular and reduced rates for the simple and complex systems. Net anterograde current is calculated as follows: (total anterograde steps - total retrograde steps)/total simulation time, across a given site averaged over 3 independent sites. Mean of 150 simulations ±95% CI of the mean is plotted across vesicle densities of 50, 100 and 200, and compared between the simple and complex systems at regular and reduced rates of motion state switching.

At the lowest densities of 50 vesicles there are hardly any vesicle clusters whose lifetimes exceed 45 seconds. At a density of 100 vesicles, the proportion of long-lived clusters increases drastically, although there are very few vesicle clusters with lifetimes longer than a minute (Fig. S3 A). In the case of 200 vesicles, the proportion of long-lived stationary vesicle clusters increases even further, with a considerable number of vesicle clusters surviving longer than a minute. This is consistent with the time-averaged distributions shown for large vesicle densities in Fig. 4, and corroborates the hypothesis that vesicle-vesicle interactions play a significant role in crowding and long-lived stationary cluster formation. In both the simple and complex systems, we see a consistent increase in proportion of long-lived stationary vesicle clusters upon a reduction in rates of motion-type switching, irrespective of the initial vesicle density (Fig. 5 B). Switching between motion states also appears to play an important role in maintaining net anterograde cargo flow along the axonal process. We find that at regular rates of switching, an increase in overall vesicle density leads to an increase in the net current (Fig. 5 C) and vesicle motion in the anterograde direction (Fig. S4) in both the simple and complex systems. The addition of obstacles such as microtubule ends and stalled mitochondria (complex system) reduces net currents across all vesicle densities tested. Although both the anterograde and retrograde currents tend to increase monotonically with increase in overall vesicle density, these increases are not linear, indicating that vesicle-vesicle interactions may interfere with directed motility, especially at higher densities. This is particularly apparent for the simple and complex systems with reduced rates, where an increase in density to 200 vesicles decreases the net current to values lower than those obtained for 50 and 100 vesicles (Fig. 5 C).

Lowering the rates of conversion from SmA to StA in systems with reduced rate reduces the number of StA vesicles compared to those in systems with regular rates. This in turn reduces the number of retrograde vesicles in this condition, as in the simulation the transitions from SmA to R motion state occur through the intermediate Step anterograde (StA) state. Thus, reducing the rates of switching between motion states also reduces the total number of reversals in the simulations. The effects of reduced rates on cargo crowding and net current (Fig. 5 B and C) appear to be largely mediated by reduced reversal rates, which suggests a role for vesicle reversals in maintaining cargo flow and easing crowding arising from vesicle-vesicle interactions.

Thus, our simulations indicate that the distribution of vesicle cluster lifetimes and net anterograde current is sensitive to overall vesicle densities as well as to the rates at which vesicles interconvert between different motion states. This has implications for cargo transport in neurons, as the stability of the cargo-motor complex may modulate vesicular motility characteristics, thereby influencing stationary cluster lifetimes and cargo flow *in vivo*.

### Tuning the locations of reversals along the neuronal process can modulate the net anterograde current of synaptic vesicles

The proportions of short-lived and long-lived stationary cargo observed in the *C. elegans* PLM neuron is best reproduced by simulations carried out for the complex system at high vesicle densities (Fig. S3 C) and reduced rates of switching between motion states (Fig. S3 B). Both these conditions have been shown to have a higher proportion of long-lived stationary clusters (Fig. 5 B and Fig. S3 C), suggesting that these neuronal processes are densely crowded with stationary vesicle clusters. However, even in the presence of such stalled vesicle clusters throughout the neuronal process, significant transport still takes place, suggesting that moving cargo employs mechanisms to navigate crowded locations. Prior studies have proposed that moving cargo can navigate obstacles by switching onto less crowded neighboring tracks or by reversing their direction of motion [34]. Additionally, results from the previous section also suggest a role for reversals in modulating cargo transport along the axonal process.

To investigate how vesicles might navigate crowded regions, we carry out simulations where both reversals and sidestepping to another microtubule track are tuned to occur at a rate that depends on the neighborhood of the cargo vesicle. Since SmA to R transitions occur through an intermediate StA state, to change reversal rates in our simulations, we alter rates of both SmA to StA and StA to R. We simulate the following limits: (1) reversals and sidestepping occur only when vesicles are stalled (at); (2) reversals and sidestepping occur anywhere along the axon, irrespective of locations of stalled vesicles and; (3) reversals and sidestepping occur only where stalled vesicles are absent (away).

We vary the ratio of reversals and side-stepping occurring ‘at’ and ‘away’ from stationary clusters, with assigned values of 100:1, 10:1, 1:1 (equal), 1:10, 1:100. In this system, currents are largest when reversals and side-stepping are allowed to occur primarily at sites of stationary vesicle clusters, as shown for ‘100:1’ (Fig. 6 A). As this ratio is decreased, net currents reduce sharply (Fig. 6 A). This variation in net current arises primarily from tuning the locations of reversals, and not from sidestepping to another microtubule track. Varying only sidestepping rates at crowded locations and keeping reversal rates constant has no effect on the net current. However, varying the rate of reversals, while keeping the sidestepping rate constant, has an effect similar to that observed when we vary both rates (Fig. 6 A). This suggests that locations at which reversals occur are critical in maintaining overall cargo flow. The simulation data additionally suggests that the net current of moving vesicles is highest when most reversals occur at stationary vesicle clusters (Fig. 6 A). However, *in vivo* we observe that reversals of pre-SVs do not occur only at regions with stationary cargo clusters (Fig. 6 B). The ratio of reversals occurring ‘at’ and ‘away’ from stationary cargo is approximately 1:1.4, similar to the 1:1 case in the simulation model (Fig. 6 B). Thus, although reversals are not triggered specifically at local roadblocks *in vivo*, they could influence overall cargo flow along the narrow, crowded geometries of the neuronal process.

**Fig 6.**
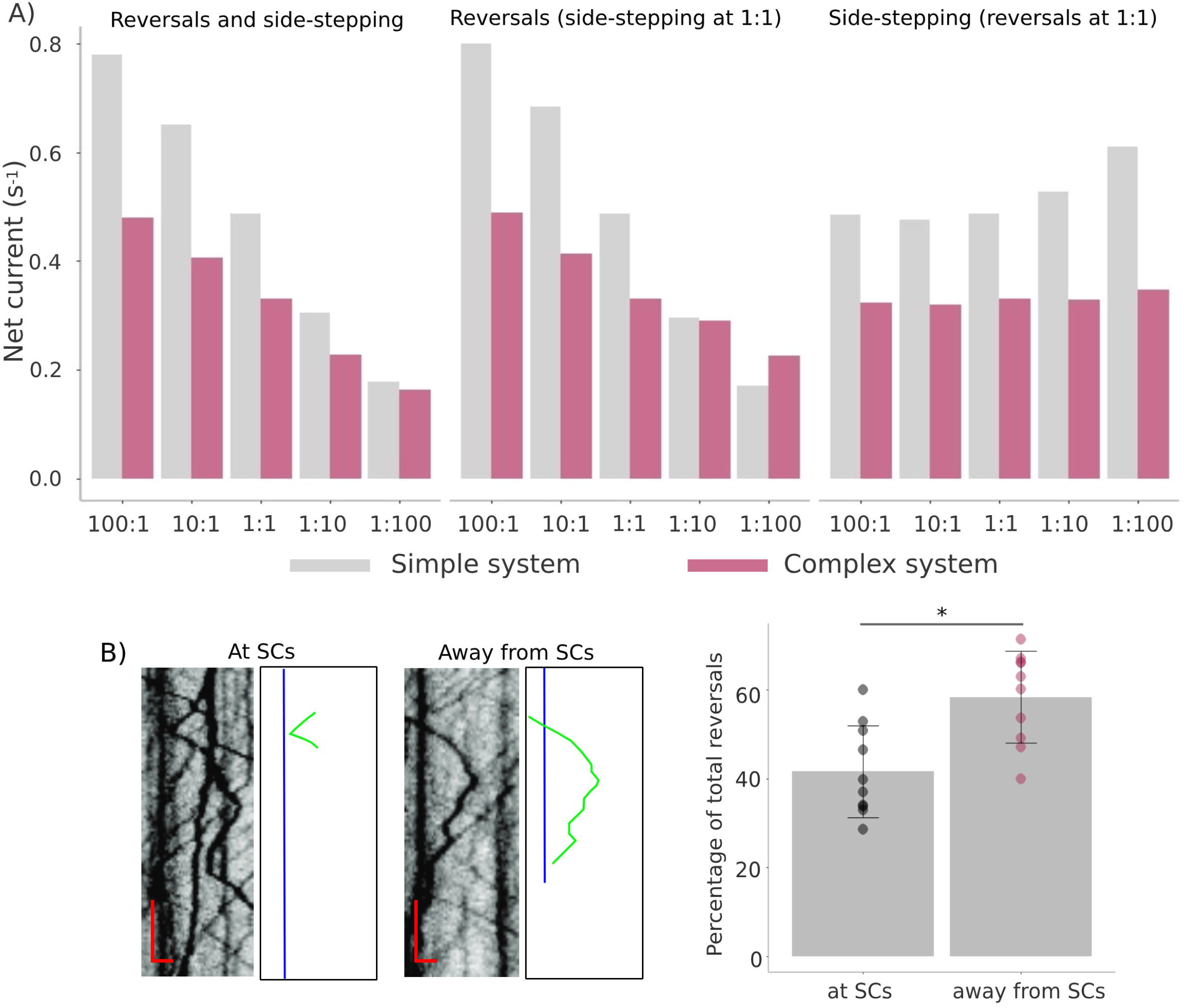
Locations of reversals influence cargo flow along the axon. (A) The panel shows a comparison between the net anterograde currents from the simple and complex systems for the following conditions-i) the rates of both reversals and sidestepping at and away from stationary vesicle clusters are varied according to the ratios shown along the x-axis, ii) only the rate of reversals at and away from stationary vesicle clusters is varied according to the ratios shown along the x-axis, while the sidestepping rate is kept constant at 1:1, iii) only the rate of sidestepping at and away from stationary vesicle clusters is varied according to the ratios shown along the x-axis, while the reversal rate is kept constant at 1:1.Data is from 150 simulations. (B) Representative kymographs depicting reversal events occurring at and away from stationary vesicle clusters in TRNs *in vivo*. Blue traces represent stationary clusters and green traces label reversing vesicles. Scale bar: x-axis = 1 *µm*, y-axis = 5sec. The plot represents the percentage of total reversals occurring at and away from stationary vesicle clusters (N = 10 animals, n (number of reversals) = 644). Mean ± SD is plotted (Student’s t-test, \**p* ≤ 0.05).

### Side-stepping and reversals are necessary to maintain net anterograde cargo transport

Since our simulations suggest that net anterograde cargo flow is sensitive to the location of occurrence of reversals and side-stepping events, we assessed the effect on cargo flux when reversals or sidestepping to another microtubule track were disallowed. Our simulations show that when sidestepping alone is disallowed, the proportion of long-lived clusters increases (Fig. 7 A and C) while the net anterograde current decreases (Fig. 7 A and D) in both the simple and complex systems. Even though the absence of sidestepping causes a comparable increase in the proportion of long-lived stationary vesicle clusters in the simple and complex systems (Fig. 7 C), the net current in the complex system is much lower as compared to the simple system (Fig. 7 D). This indicates that sidestepping to another microtubule track plays an important role in vesicles navigating the crowded locations created by long-lived stationary vesicle clusters. We infer that systems which are densely crowded with stationary vesicle clusters, like the TRNs of *C. elegans*, may rely more on the ability of vesicles to sidestep onto neighboring microtubules to maintain cargo flow, as compared to less crowded systems.

**Fig 7.**
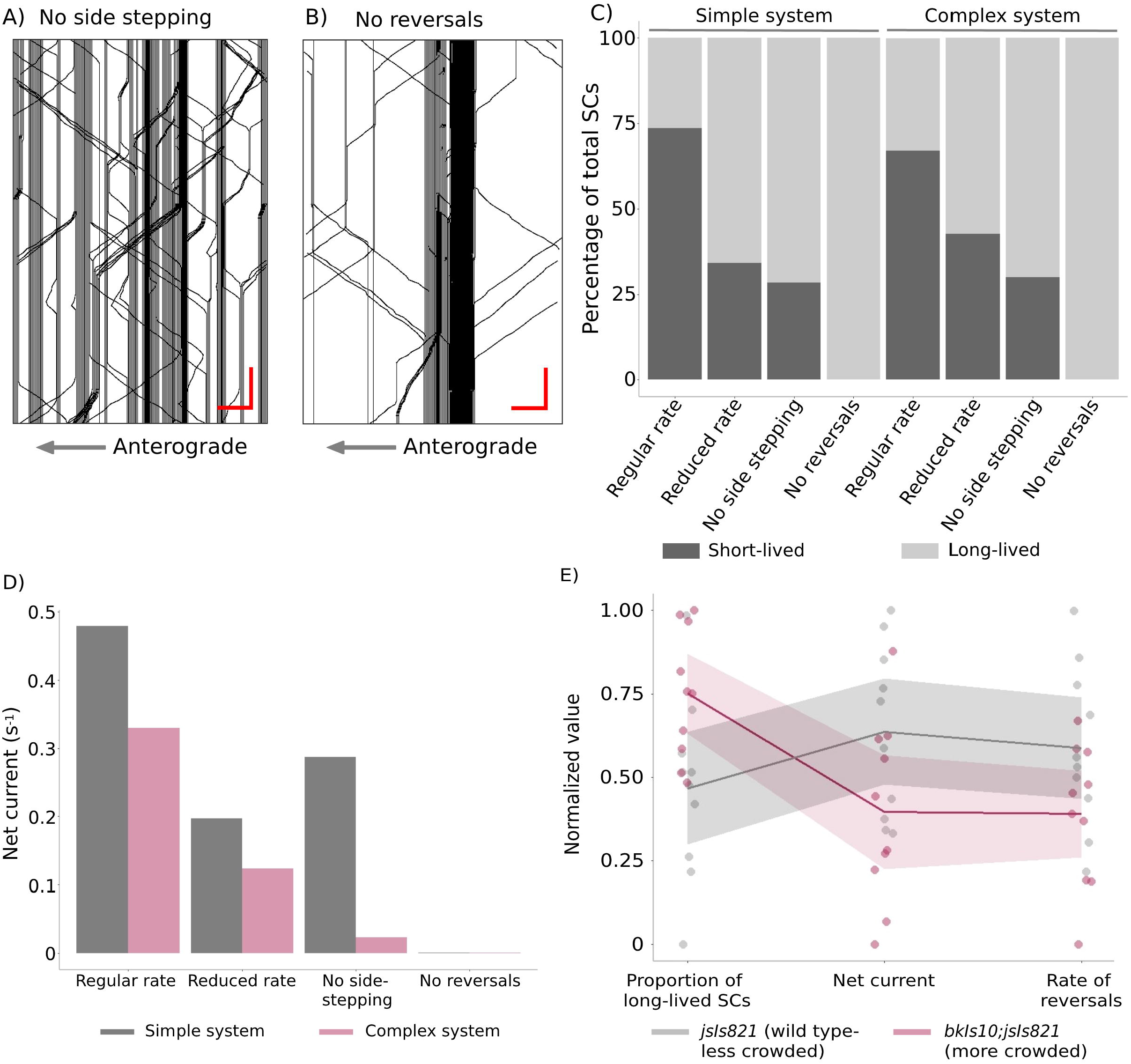
Reversals and side-stepping are important for reducing cargo crowding and maintaining flux along the axon. (A, B) The kymographs respectively depict vesicle transport predicted by simulations when no sidestepping or reversals are allowed. In the absence of reversals we show the kymograph before the final state, a single infinitely long-lived cluster, is attained. Scale bar: x axis = 0.5 *µm*, y axis = 2 sec. (C) The plot shows the proportions of total stationary clusters that are short-lived and long-lived across different simulation conditions. Stationary cluster numbers are pooled across 150 simulations for each condition. (D) The panel shows net anterograde current calculated as follows: (total anterograde steps - total retrograde steps)/total simulation time, across a given site averaged over 3 independent sites and 150 simulations. (E) The panel depicts a parallel plot that compares normalized values of the proportion of long-lived SCs, net current, and reversal rate between wild type (*jsIs821*) and the tauopathy model (*bkIs10*;*jsIs821*) in 5-day adult *C. elegans* hermaphrodites. The normalization and plotting is described in detail in the ‘Methods’ section (“Normalization of data for parallel plots”). The solid circles represent data points from individual animals, the solid lines represent mean values for each category, and the shaded bands represent the 95%CI around the mean. N = 10 animals for each genotype, n(*jsIs821*) = 3064 moving vesicles, 277 stationary vesicles, n(*bkIs10*;*jsIs821*) = 1958 moving vesicles, 370 stationary vesicles.

In the absence of reversals, the system reaches a final state where all currents vanish and a single large stationary vesicle cluster dominates (Movie S1, Fig. 7 B-D). Starting from such a jammed configuration, allowing for vesicle reversals dissolves the block, leading to a net current (Movie S2). This is consistent with the idea that allowing for reversals prevents vesicular crowding [34, 35]. Our simulation suggests that (i) the ability of moving vesicles to reverse is crucial to avoiding a permanently blocked state with no current and that (ii) side-stepping plays an important, but secondary, role in maintaining cargo flow along crowded neuronal processes.

In order to validate the predictions from our simulations, we examined a tauopathy model of *C. elegans, bkIs10*, which expresses the IN4R isoform of human tau carrying the V337M FTDP-17 (Frontotemporal Dementia with parkinsonism chromosome 17 type) mutation. *C. elegans* with *bkIs10* form Tau aggregates along the neuronal process, have uncoordinated movements, and show age-dependent loss of axons [36]. We find that transport of pre-SVs in *bkIs10* animals is comparable to transport in wild type animals at younger stages (L4 and 1-day) (Fig. S5 A-D). In 3-day and 5-day adults *bkIs10* animals have a significantly higher density of stationary vesicle clusters with a greater proportion of long-lived stationary vesicles clusters compared to wild type (Fig. S5 A, B, and F, Fig. 7 E). Thus, the tauopathy model at older stages presents a system that is more densely crowded with stationary vesicle clusters than wild type, which can be used to investigate the correlation between reversal rate, net current, and the proportion of long-lived stationary clusters *in vivo*. The phenomenon of side-stepping is not observable along the narrow processes of *C. elegans* neurons due to the technical limitations of existing live imaging techniques. We observed that the reversal rate in *bkIs10* animals was comparable to that of wild type animals at younger stages (Fig. S5 C and E), but was significantly reduced in 3-day and 5-day old animals (Fig. S5 C and F, Fig. 7 E). Interestingly, we found that the net current in *bkIs10* animals was comparable to wild type animals from L4 to 3-day adults (Fig. S5 D-F), and was only significantly reduced in 5-day adults (Fig. S5 D, Fig. 7 E). This suggests that a decrease in reversal rate precedes the decrease in net current observed along the neuronal process (Fig. S5 D and E).

Our experimental and simulation data collectively suggest that a system that permits reversals and side-stepping, irrespective of their location, is sufficient to maintain a steady anterograde current of synaptic vesicles along the neuronal process (Fig. 6 A, Fig. 7). We find that the proportions of long-lived stationary vesicle clusters is negatively correlated to net cargo flow and reversal rate (Fig. 7 C-E), indicating that stationary cargo clusters might effectively function as reservoirs, holding and releasing cargo so that, on average, a steady flux is maintained along the axon. This suggests that the previously proposed reservoir hypothesis [8] underlies the physiological significance of stationary synaptic vesicle clusters and that motion state changes could play an important role in modulating the dynamics and lifetimes of such clusters.

## Discussion

The narrow and highly confined processes of *C. elegans* TRNs present specific obstacles to transport. Stationary vesicle clusters, which are a prominent feature of axonal transport *in vivo*, are crowded locations [8]. At these locations, moving vesicles stall for timescales ranging from a few seconds to several minutes. Since crowded locations pose impediments to transport by causing vesicle accumulation, potentially choking off steady vesicular flux, it appears reasonable that neurons may have evolved strategies to cope with the consequences of crowding in a dense environment. Indeed, large stationary accumulations of vesicles and low vesicular flux are prominent features of a number of neurodegenerative diseases. This suggests that understanding the *in vivo* mechanisms leading to the formation and dissolution of such stationary vesicle clusters may provide insights into such disease states [37, 38].

Prior to studying specific disease states, it is important to examine how vesicle clusters might form and persist for a broad range of times in healthy neuronal processes, while still allowing for steady cargo flux. It has been previously reported that stationary vesicle clusters, although long-lived, are dynamic, dissolving through the same microscopic processes that led to their formation [8]. While detachment from microtubules and diffusion is one possible solution to bypass crowded locations, diffusive processes are expected to play a subordinate role in cytoskeletally dense neurons. This is evident in our FRAP experiments conducted in *C. elegans* PLM neurons, wherein all the fluorescence signal recovery observed in the bleached region over a timescale of minutes is associated with active vesicular transport (Fig. 2 A and H).

We constructed a simulation model for the axonal transport of pre-synaptic vesicles that was benchmarked against the experimentally-derived ratio of motion states and average velocities observed for moving precursors of synaptic vesicles in *C. elegans* touch receptor neurons. Novel *in silico* methods were developed to simulate photobleaching and axotomy. We further systematically varied multiple simulation parameters, narrowing them down to a set of conditions that produced results consistent with experimental features.

Our simulations indicate that the transport characteristics of synaptic vesicles play a hitherto unappreciated role in maintaining stationary cluster lifetimes and net cargo flow along the neuronal process. The dynamics of stationary vesicle clusters is sensitive to the rates at which cargos switch between three experimentally observed motion states. Altering these rates leads to changes in vesicle cluster sizes, vesicle cluster lifetimes and net cargo flow (Fig. 5). The ability to tune rates of conversion, perhaps by activating or inactivating a class of motors consequent to a signalling event *in vivo*, could potentially enhance or suppress regular cargo transport [25, 39, 40]. We further conclude that permanent traffic jams are avoided because such clusters are dynamic, and their lifetimes are governed by vesicle motility characteristics.

Two such motility characteristics that could regulate cargo crowding are sidestepping to another microtubule track and reversals. Several biophysical and modelling studies have proposed that moving cargo can navigate crowded locations through track switching, enabled by the association of cargo-bound motors with multiple microtubules [34]. Sidestepping provides a means of spatial relaxation of crowding, allowing vesicles to switch from more crowded to less crowded microtubule tracks, thereby maintaining cargo flow. Our simulations show that preventing vesicles from sidestepping onto neighboring microtubules leads to a significant reduction in vesicular transport and a concomitant increase in stationary cluster formation along the neuronal process (Fig. 7 C and D).

Reversals may also aid in avoiding crowded environments by allowing moving cargo to sample multiple microtubules, and redistributing cargo to avoid pile-ups at crowded locations [34, 35]. Reversals can reduce the frequency of vesicle-vesicle interactions at crowded locations by causing cargo to change their direction of motion. A temporal relaxation of crowding may be achieved when a vesicle that has undergone reversal returns to the same region of the axon or microtubule track at a later time. Our simulations suggest that spatial relaxation of crowding through sidestepping plays an important role in maintaining cargo flow in neurons. However, in vesicle-dense, crowded environments, reversals may play a more important role in maintaining cargo transport (Fig. 6 A, Fig. 7 C, D).

On assessing motion properties of pre-SVs in a neurodegeneration model of *C. elegans* [36], we observed that an increase in stationary cargo density with age was accompanied by a decrease in reversal rate in this diseased condition (Fig. 7 E, Fig. S5 A-C). This decrease in reversals and increase in stationary cargo was consequently observed to affect net cargo flow as predicted by our simulation model (Fig. 7 D and E, Fig. S5 D). Additionally, wild type animals independently showed a decline in stationary cargo density with age and a corresponding increase in reversal rate (Fig. S5 A-C). This age dependent transition observed in wild type animals was completely lost in the tauopathy model (Fig. S5 A-C). The absence of such an age dependent modulation in reversal rate in the tauopathy model may contribute to crowding and influence disease progression [34, 37, 38, 41, 42].

Collectively, our experimental observations and simulations predict that a reduction in the ability of vesicles to i) switch between distinct motion states, ii) sidestep onto different microtubules during transport, or iii) reverse their direction of transport, leads to increased cargo stalling and a decrease in net anterograde flux (Fig. 7). All these are characteristic features of the defective axonal transport observed in several neurodegenerative disease conditions [37, 41, 43–49]. This suggests that the age-dependent decline in axonal transport in neurodegenerative disease models could be driven by changes in vesicle motility characteristics. Studying these changes may provide a new window into a class of neurodegenerative diseases and their associated therapeutics

## Methods

### Worm strains and maintenance

The strains used in this study are listed in the Table S1

All *C. elegans* strains used in this study were maintained at 20*°* C on Nutrient Growth Media (NGM) media seeded with *E. coli* OP50, as described in the standard protocol [50]. The animals were primarily imaged at the L4 stage. In some experiments, animals were imaged at the 1-day, 3-day and 5-day adult stages. This is mentioned in the respective sections. All animals used for imaging came from non-contaminated, growing plates.

### Time-Lapse Imaging

Live single worms were mounted on glass slides with 5% agar pads, and anesthetized using 5mM Tetramisole (Sigma-Aldrich, St. Louis, MO, USA) prepared in the M9 buffer. All time lapse imaging was conducted on the Yokogawa CSU-X1-A3 spinning disc, using the Hamamatsu ImagEM C9100-13/14 EMCCD Camera integrated with an Olympus IX83 microscope by Perkin Elmer. Samples were imaged using a 100X 1.63N.A. objective. Some imaging was also done on the Zeiss LSM 880 using the 63X 1.4N.A. objective. Imaging was done at 5fps and 3-5min movies were taken for all genotypes.

### Image Analysis

All image panels used for representation and analysis of time lapse movies were generated using Fiji-ImageJ v1.52p. Experimental kymographs were generated using the MultipleKymograph plugin. Plugins were downloaded from the NIH website with the following links;

http://www.rsbweb.nih.gov/ij/ and http://www.emblheidelberg.de/eamnet/html/bodykymograph.html.

In the kymograph, cargo moving in the retrograde direction (towards the cell body), and anterograde direction (away from the cell body) appear as sloped lines, while stationary cargo appear as vertical lines. A cargo is counted as moving if it has been displaced by at least 3 pixels in successive time frames.

### Calculation of motion proportions (experiment)

Moving vesicles were manually annotated on experimental kymographs, with individual trajectories traced as accurately as possible, using the ‘Segmented Line’ feature in Fiji. Motion parameters such as run length and pause duration were calculated using an in-house macro (‘macro for motion properties.ijm’) written in Fiji. The code for the same can be found on the following GitHub repository link-https://github.com/amruta2612/Fiji-macros. Segment run length is the distance moved by the cargo between pauses, using a conversion factor of 0.13 *µm*/pixel. Pause time is defined as the length of time for which a cargo stays stationary between two consecutive runs. The ‘Measure’ function was used to obtain the length of these lines in pixels, which was then converted to units of time using a conversion factor of 1 pixel= 0.2s (corresponding to 5fps). Total run length is defined as the sum of segment run lengths along the trajectory of individual vesicles.

### Stationary cargo analysis (experiment)

All vertical lines on the kymograph represent stationary/paused cargo. The entire length for which each particle remains stationary was annotated using the Segmented Line feature in Fiji (ImageJ). The length of each line was calculated in pixels using the measure tool of ImageJ. This was used to calculate the time for which the vesicle remains stationary (in seconds) based on the frame rate at which the movie was captured. A vesicle was counted as stationary if it did not move for more than 15seconds (for synaptic vesicles). This cut off is based on the average pause times of moving vesicles and any vesicle is said to be stationary if it remains paused for at least five times longer than its average pause time. All stationary vesicles were further binned as short-lived or long-lived based on the time duration for which it remained stationary. For this, we tracked two types of stationary vesicles in our kymographs, 1) stationary vesicles whose end and origin was within the time window of the entire kymograph analysed, 2) a greater than category (>) which includes stationary vesicles whose end or origin was not within the time window of the kymograph analysed, and thus its exact duration was not known. All vesicles under type (1) that remained stationary for 15-45 seconds (Fig. S1 F) were binned into the short-lived category and those beyond 45 seconds (45s-1min, 1-5mins in Fig. S1 F) were binned into the long-lived category. All vesicles under type (2), the greater than (>) category (> 15*s*, > 45*s*, > 1min in Fig. S1 F), were binned into the long-lived category as long as it met the minimum cut-off of 15seconds. This was done as we do not know the exact time duration for which these vesicles are stationary. All bins are non-overlapping (Fig. S1 F). All stationary cargo were annotated manually and were counted and binned into the relevant categories using a macro code (‘macro for SC lifetime.ijm’) written in ImageJ. The code for the same can be found on the following GitHub repository link-https://github.com/amruta2612/Fiji-macros.

### Analysis of stationary cargo site occupancy (experiment)

Stationary cargo were identified as described in the previous section. In the kymograph, we marked each stationary cluster using a rectangular ROI, with the length representing the lifetime of the stationary cluster (in secs), and the width representing the distance occupied by the stationary cluster along the axonal process (in microns). The area of this rectangular ROI thereby represents the total distance along the axon that is occupied by the respective stationary cargo over time. In order to quantify and compare stationary cargo site occupancy over time, the sum of the areas of all rectangular ROIs was plotted as a percentage of the total area of the kymograph.

### Flux analysis (experiment)

Sloped lines in the kymograph indicate moving vesicles. The minimum cut-off used for counting a sloped line as a moving cargo was a displacement of 3 pixels in successive time frames. All vesicles moving away from the cell body were counted as anterograde flux, while the ones moving towards the cell body were counted as retrograde flux. Total flux was calculated as the sum of anterograde and retrograde flux, while net current was calculated as the difference between anterograde and retrograde flux normalized to time, expressed as number/(*µm*\*min). All moving vesicles were annotated manually and were counted and binned into the relevant categories using a macro code (‘macro for flux.ijm’) written in ImageJ. The code for the same can be found on the following GitHub repository link-https://github.com/amruta2612/Fiji-macros.

### Reversal analysis (experiment)

Reversals refer to changes in the direction of motion of moving vesicles from either anterograde to retrograde direction or vice versa. An anterogradely moving vesicle that changed its direction to retrograde along its trajectory was counted as an anterograde reversal. Similarly, any vesicle moving in the retrograde direction switching to the anterograde direction was counted as a retrograde reversal. Reversal data for all experiments is represented as a fraction of moving vesicles in a particular movie, i.e., the number of reversals per 10 moving vesicles. For anterograde reversal rate calculation, anterograde reversals were divided by anterograde flux; for retrograde reversal rate calculation, retrograde reversals were divided by retrograde flux and for total reversal rate, total reversals were divided by total flux.

### Locations of reversals (experiment)

The locations of reversals were divided into two categories: 1) at sites of stationary vesicles: all reversal events that occur on encountering a stationary vesicle or within the proximity of 2 pixels (0.26 *µm*) on either side of stationary vesicles were counted in this category and 2) away from stationary vesicles: all reversal events that occur at a distance of > 2pixels (0.26 *µm*) from stationary vesicles were counted in this category. This data was then represented as the percentage of reversals taking place at or away from a stationary cluster. Since stationary vesicles are bright, these cut-offs were chosen keeping in mind the diffraction limits of microscope and accuracy of measuring distance from stationary vesicles.

### Fluorescence recovery after photobleaching (experiment)

Photobleaching experiments were performed on an Olympus FV3000 confocal microscope using a 40X/1.3N.A. oil objective, where a 10 *µm* region of PLM neuronal process near the cell body was imaged using a 488nm laser for 1 minute before bleaching, and for 3 minutes immediately after bleaching, to monitor fluorescence recovery over time. The analysis of bleach recovery was conducted as follows-In each movie, the 10 *µm* region that was photobleached was marked as the ‘bleached region’, a 10 *µm* region excluding the neuronal process (chosen arbitrarily) was selected as the background, and a 10 *µm* region of the neuronal process excluding the bleached region (chosen arbitrarily) was selected as the ‘unbleached region’. The mean fluorescence intensity across each 10 *µm* region was monitored over time. The normalized intensity for each timepoint was calculated as follows-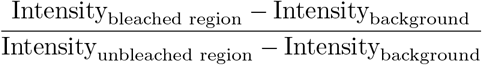

### Axotomy (experiment)

Axotomy experiments were performed using a 355nm (UV) pulsed nanosecond laser (Minilite Series, Flashlamp pumped, Q-Switched, Nd YAG) operated in low energy mode at a repetition rate of 10. Time-lapse fluorescence images of specific regions of the neuronal processes of TRNs were acquired at 5fps, using a 100X/1.4 NA oil objective on the Olympus IX83 microscope integrated with the Yokogawa CSU-X1-A3 spinning disc and Hamamatsu ImagEM C9100-13/14 EMCCD Camera (by Perkin Elmer). The analysis for vesicle accumulation following axotomy was conducted as follows-

For each animal/movie, the fluorescence intensity over a 1 *µm* region at the proximal and distal cut sites was monitored at 1min20s, 2min20s, and 3min20s post-axotomy, using the ‘Segmented Line’ and ‘Plot Profile’ features in Fiji. At each timepoint, 6 independent background measurements were taken (1 *µm* each) and averaged to obtain the background intensity value for each timepoint post-axotomy. The normalized intensity values over a 1 *µm* region for each timepoint were calculated using the formula

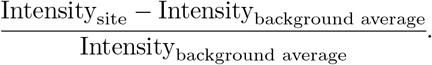

The normalized intensity values were averaged across animals and the average normalized intensity values were fitted to an exponential decay curve as follows-

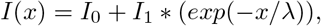

where I(x) represents the intensity at a distance of ‘x’ *µm* from the cut site, *I*_0_ represents the baseline intensity value, *I*_1_ represents the scaling factor, and λ is the decay constant.

### Normalization of experimental data for parallel plots

For each categorical variable, such as ‘Proportion of long-lived SCs’, ‘Net current’, and ‘Rate of reversals’, all data points from both wild type (*jsIs821*) and the tauopathy model (*bkIs10;jsIs821*) were pooled together and sorted in ascending order of values. The maximum and minimum values were noted and each data point was transformed using min-max normalization: Transformed value= (Original value - Minimum value)/(Maximum value - Minimum value). All the transformed data points now lie between 0 and 1. This transformation was applied independently to data for each categorical variable, and the final transformed data points were color-coded according to their respective genotypes.

### Monte Carlo Simulations

For most of our simulations, excluding our results for simulated photobleaching and axotomy, we consider a system of 10 microtubules, each containing 1000 sites, representing motor binding sites on parallel tracks with lattice points spaced 8nm apart from each other. Since properties of current and stationary clusters are independent of system size for a settled system, simulations with smaller system sizes - that reduce the time taken to run the simulation - suffice. Larger system sizes (a system of 10 microtubules each containing 8000 sites, thus simulating an 64 *µm* section of the axon) were used for photobleaching and axotomy simulations. The numbers of vesicles are scaled accordingly so that the same range of densities is sampled in both the small and the large system. The use of the larger system in the photobleaching simulation is required so that the re-entry of bleached vesicles, a possibility with periodic boundary conditions, can be ruled out.

In all simulations, vesicles, at any given time, are associated with a single lattice point and hop at fixed rates between neighboring lattice points. Three different motion states of vesicles are modeled based on experimental data: i) smooth anterograde, ii) step anterograde and iii) retrograde. Vesicles are assumed to exclude other vesicles from a region of two lattice points on either side, given the (approximately two times larger) size of the cargo in comparison to that of the motor. Hopping rates for the three types of vesicles are chosen to represent the experimental rates, consistent with measured properties of isolated vesicles away from the locations of stationary vesicle clusters.

Apart from moving one site at a time in a preferred direction a vesicle can: a) the vesicle can move to an adjacent site on another track (side-stepping) at fixed rate, b) it can change state, switching between SmA, StA and R states at a prescribed rate (see Table S2). Vesicles can be blocked from moving, either by another vesicle, a microtubule end or a large obstacle such as a mitochondrion. We consider several limits: i) vesicles can sidestep or change state irrespective of whether they are blocked ii) vesicles can sidestep or change state only when they are not blocked and iii) vesicles can sidestep or change state only when they are blocked. In simulations, the motion path of each vesicle is tracked and parameters like velocity, pause time, pause frequency, run length, and flux are calculated. For further details, see supplementary methods. The choice of rates is discussed in more detail in Supplementary Information.

### Simulation Kymographs

Since all vesicles are tracked in the simulations, it is easy to generate kymograph tracks from them. In order to make these tracks visually similar to experimental kymographs, we assign different grey-scale values and line thicknesses to vesicles depending on their motion. With this in mind, we choose the following algorithm for generating our kymographs. We plot all vesicles, with the structure of the kymograph representing the relative proportion of stationary vs non-stationary vesicles. In our kymographs, there are 2 parameters for the tracks, the greyscale value and the thickness. Of all the vesicles, we identify those who are part of long-lived clusters, representing them by thicker (darker) lines in the kymograph. The grey-scale is assigned in proportion to the time spent by such vesicles in stationary clusters. To set these (thickness and greyscale) values, we compute the maximum time each vesicle spends at a site, assigning a line thickness proportional to the time spent in a stationary state. For the greyscale value, a lighter shade of grey is used for vesicles that show substantial movement, whereas we use a darker shade of grey (RGB value [0.1, 0.1, 0.1] for vesicles that are largely stationary. These choices yield simulation kymographs that visually resemble those obtained from experiments.

### Computing stationary vesicle cluster lifetimes in simulation

We take simulation snapshots of the axon at time intervals which correspond to the frame rate of experimental movies. We collapse sites with the same index across microtubules, since vesicles on different tracks corresponding to the same indexed site cannot be resolved separately in experiments. A site is considered to be occupied, contributing to the fluorescence signal, if the corresponding site in any of the microtubules is occupied. When a site is occupied across two adjacent snapshots, we consider it to be a stationary cluster and the duration of the respective stationary vesicle cluster is incremented by an amount equivalent to the time interval of the snapshot. The duration assigned to a stationary vesicle cluster is computed as the time duration for which the stationary vesicle cluster is observed within the time window of the simulation, irrespective of whether the start or end point of the cluster lies completely within the time window of the simulation snapshot.

### Estimating the number of long-lived stationary clusters per 8s microns in simulation

As in the experiments, long-lived stationary vesicle clusters are defined as those clusters that are observed to survive for longer than 45s. But in the simulation, all stationary vesicle clusters lasting longer than 45s are counted irrespective of whether their start or end points are completely within the time window of the simulation.

### Calculating fraction of axon occupied by all stationary vesicle clusters in simulations

Stationary cargo site occupancy analysis for the simulation data is performed in a manner similar to that in the experiment. The rectangular ROI for each stationary cluster in the kymograph is calculated as the product of the lattice width and the time length occupied by the stationary cluster. The rectangular ROI data for all stationary clusters in the system is then represented as the fraction of the total area of the kymograph.

### Estimating current in simulations

The current (equivalently the flux) is obtained by computing the time-averaged currents of different states of vesicles in the following way: Choosing a link connecting two neighbouring sites, we compute the number of anterograde vesicles of both states (smooth and step) moving across it, as well as the number of retrograde vesicles, over an interval of time. Subtracting the numbers of retrograde vesicles from anterograde vesicles, and dividing by the elapsed time, gives us the current, which can then be computed over a large number of configurations to generate averages in steady state.

### Simulating Fluorescence Recovery After Photobleaching

Fluorescence Recovery After Photobleaching (FRAP) experiments are simulated on a 64 *µm* lattice with 400, 800 and 1000 vesicles. Once the system settles into a steady state, vesicles in a 8 *µm* section of the lattice are tagging as ‘bleached’ - in effect hiding them from that point on in the simulation. The recovery of unbleached vesicles in the bleached region is then followed over time. Though the bleached vesicles continue to hop, change track, change type and interact as before, they are excluded from subsequent analysis of bleach recovery and generation of kymographs.

### Simulating axotomy

We simulate axotomy by blocking a group of 100 sites (0.8 *µm*) on all contiguous microtubules. Vesicles can thus move up to but not across the axotomized region. This leads to an accumulation of vesicles over time and subsequent saturation of the density at and close to the two extremal sites of the axotomized region. We plot averaged densities of vesicles of all motion states in the vicinity of the axotomized region, thus providing predictions for the experiments.

## Acknowledgments

IMSc for computational facilities, PRISM project for funding (VK)

