## Supplementary Figures for "Cargo crowding, stationary clusters and dynamical reservoirs in axonal transport"

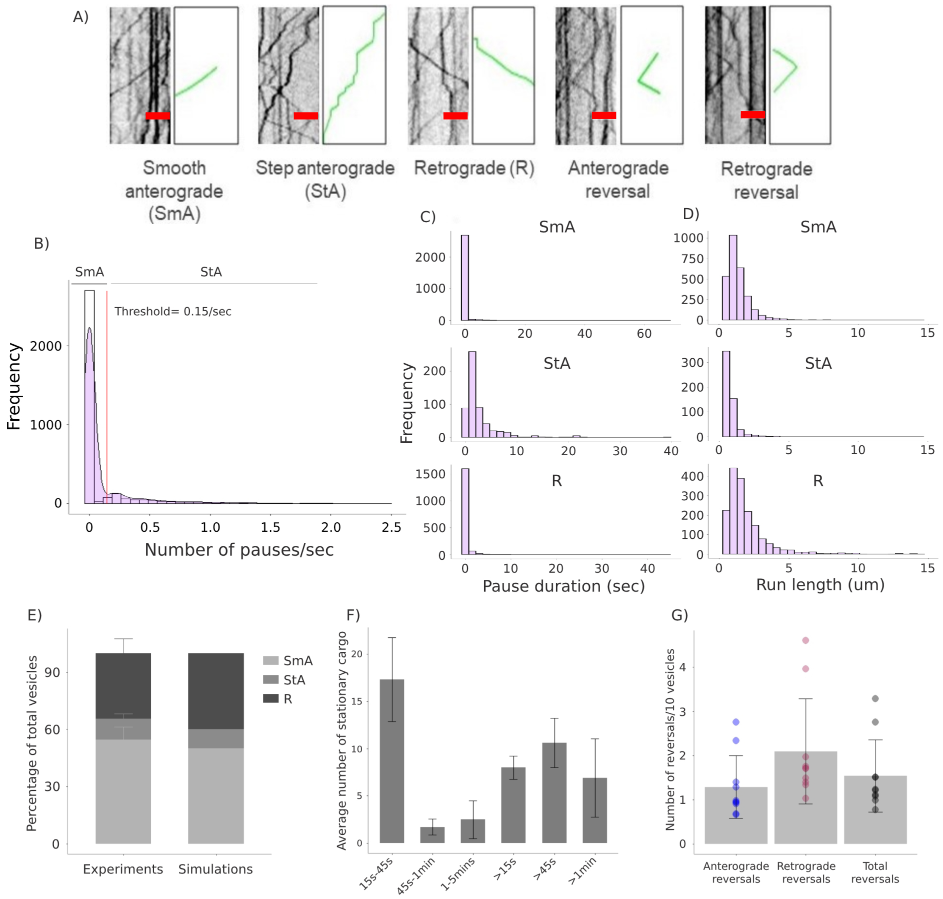
A)
Smooth anterograde (SmA)
Step anterograde (StA)
Retrograde (R)
Anterograde reversal
Retrograde reversal
B)
SmA
StA
Threshold= 0.15/sec
Frequency
Number of pauses/sec
C)
SmA
StA
R
Frequency
Pause duration (sec)
D)
SmA
StA
R
Run length (um)
E)
Percentage of total vesicles
SmA
StA
R
Experiments
Simulations
F)
Average number of stationary cargo
15s-45s
45s-1min
1-5mins
>15s
>45s
>1min
G)
Number of reversals/10 vesicles
Anterograde reversals
Retrograde reversals
Total reversals

**Figure S1**

**Figure S1. Characterization of general features of synaptic vesicle transport *in vivo***

(A) Representative kymographs of GFP::RAB-3 marked pre-SVs, showing examples of smooth anterograde (SmA), step anterograde (StA), retrograde (R) motion states, and anterograde and retrograde reversals. Scale bar: x-axis = 3 $\mu m$. All representative kymographs have a total time duration of 16secs.

(B) The plot shows a histogram of the number of pauses exhibited by anterogradely moving vesicles along their trajectory, normalized to time. The data is pooled across 10 animals, with a total of 3310 anterograde vesicles analyzed.

(C, D) The plots show histograms of pause duration (in seconds) and run length (in $\mu m$) respectively, for vesicles exhibiting different motion states. The data is pooled across 10 animals. Total number of SmA vesicles = 2755, StA = 555, and R = 1695.

(E) The plot shows the relative proportions of the three vesicle motion states observed in experiments. The data is calculated across 10 animals, with Mean $\pm$ SD plotted for each category. Simulations are benchmarked to the ratio of motion states observed in experiments, which is 50 (SmA):10 (StA):40 (R).

(F) The plot represents the average number of stationary vesicle clusters observed for different lifetimes. N = 10 animals, n = 378 stationary vesicles, with Mean $\pm$ SD plotted for each category.

(G) The plot shows anterograde, retrograde and total reversals per 10 moving vesicles of GFP::RAB-3 marked pre-SVs (reversal rate) . N = 10 animals, n = 4794 moving vesicles, with Mean $\pm$ SD plotted. One-way ANOVA, with Bonferroni post hoc test was performed, and none of the categories were found to be significantly different from each other.

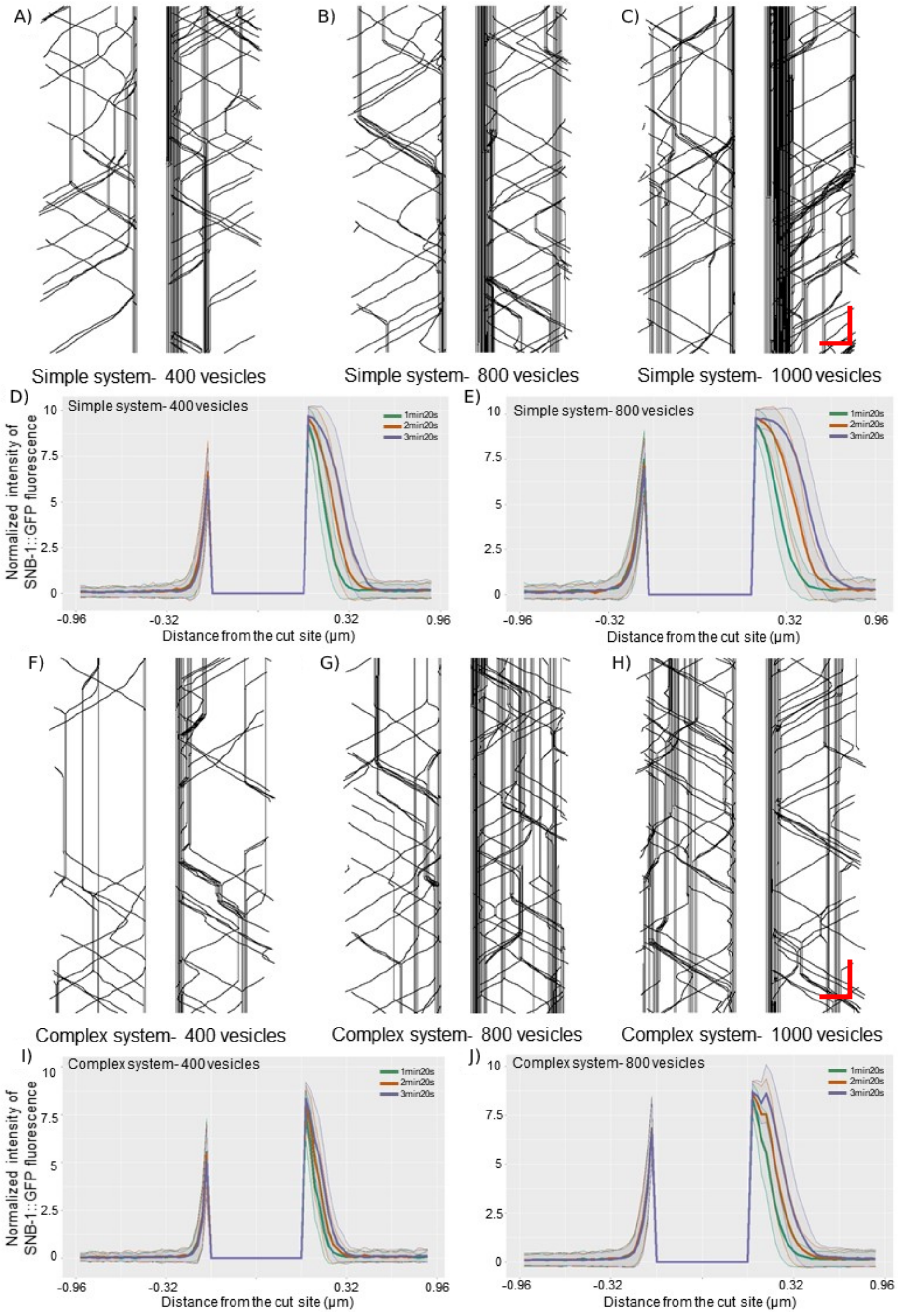

A)
B)
C)
Simple system- 400 vesicles
Simple system- 800 vesicles
Simple system- 1000 vesicles
D)
Simple system- 400 vesicles
E)
Simple system- 800 vesicles
1min20s
2min20s
3min20s
Normalized intensity of SNB-1::GFP fluorescence
Distance from the cut site (µm)
F)
G)
H)
Complex system- 400 vesicles
Complex system- 800 vesicles
Complex system- 1000 vesicles
I)
Complex system- 400 vesicles
J)
Complex system- 800 vesicles


**Figure S2**

**Figure S2. Simulations reproduce vesicle accumulation profiles observed after axotomy *in vivo***
(A-C) & (F-H) Representative kymographs showing vesicle transport and accumulation as predicted by different simulation conditions, 3min20s post-axotomy. All kymographs are oriented such that the anterograde direction is from right to left, with the proximal cut site to the right of the axotomized region and the distal cut site to the left. Scale bar: x axis = 0.5 $\mu m$, y axis = 2 sec.
(D) The panel shows the normalized mean density profiles of vesicles at 1min20s, 2min20s and 3min20s post-axotomy in the simple system with an initial vesicle density of 400 vesicles, with the standard deviations for each shown as grey bands. Data from 208 simulations.
(E) The panel shows the normalized mean density profiles of vesicles at 1min20s, 2min20s and 3min20s post-axotomy in the simple system with an initial vesicle density of 800 vesicles, with the standard deviations for each shown as grey bands. Data from 137 simulations.
(I) The panel shows the normalized mean density profiles of vesicles at 1min20s, 2min20s and 3min20s post-axotomy in the complex system with an initial vesicle density of 400 vesicles, with the standard deviations for each shown as grey bands. Data from 161 simulations.
(J) The panel shows the normalized mean density profiles of vesicles at 1min20s, 2min20s and 3min20s post-axotomy in the complex system with an initial vesicle density of 800 vesicles, with the standard deviations for each shown as grey bands. Data from 494 simulations.

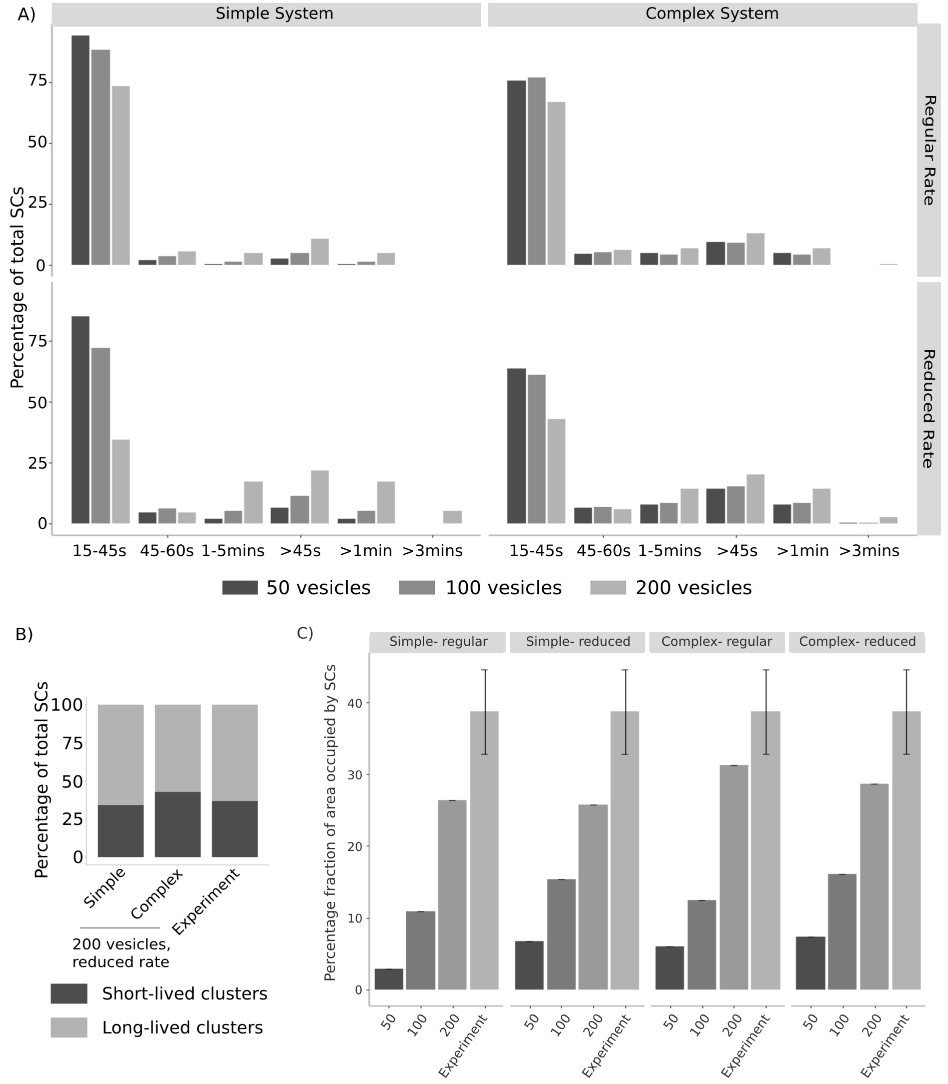
A)
Simple System
Complex System
Regular Rate
Reduced Rate
Percentage of total SCs
0
25
50
75
15-45s
45-60s
1-5mins
>45s
>1min
>3mins
50 vesicles
100 vesicles
200 vesicles
B)
Percentage of total SCs
0
25
50
75
100
Simple
Complex
Experiment
200 vesicles, reduced rate
Short-lived clusters
Long-lived clusters
C)
Simple- regular
Simple- reduced
Complex- regular
Complex- reduced
Percentage fraction of area occupied by SCs
0
10
20
30
40
50
100
200
Experiment

**Figure S3**

**Figure S3. Complex system reproduces the lifetimes of stationary vesicle clusters observed *in vivo***

(A) The panel shows the distribution of lifetimes of stationary vesicle clusters observed in our simulations, across different vesicle densities, at regular and reduced rates for the simple and complex systems. Total number of stationary vesicle clusters are pooled across 150 simulations for each condition, and each simulation condition is compared to experiments (data pooled across 10 *jsIs821* animals). Add total no of SCs analyzed.

(B) The panel compares the proportions of short-lived and long-lived stationary vesicle clusters observed in the simple and complex systems at reduced rates and a vesicle density of 200, with experiments. The simulation data for the simple system has N = 150 simulations, nSL (number of short-lived stationary vesicles) = 13344, nLL (number of long-lived stationary vesicles) = 25630. The complex system has N = 150 simulations, nSL = 26922, nLL = 36059. For the data from experiments, N = 10 *jsIs821* animals, nSL = 173, nLL = 297.

(C) The panel compares the area of kymograph occupied by stationary vesicle clusters over time across various simulation conditions with experimental data. Data for the simple and complex systems is pooled across 150 simulations. For the experimental data N = 10 animals, n = 470 stationary vesicles. Mean $\pm$ SD plotted.

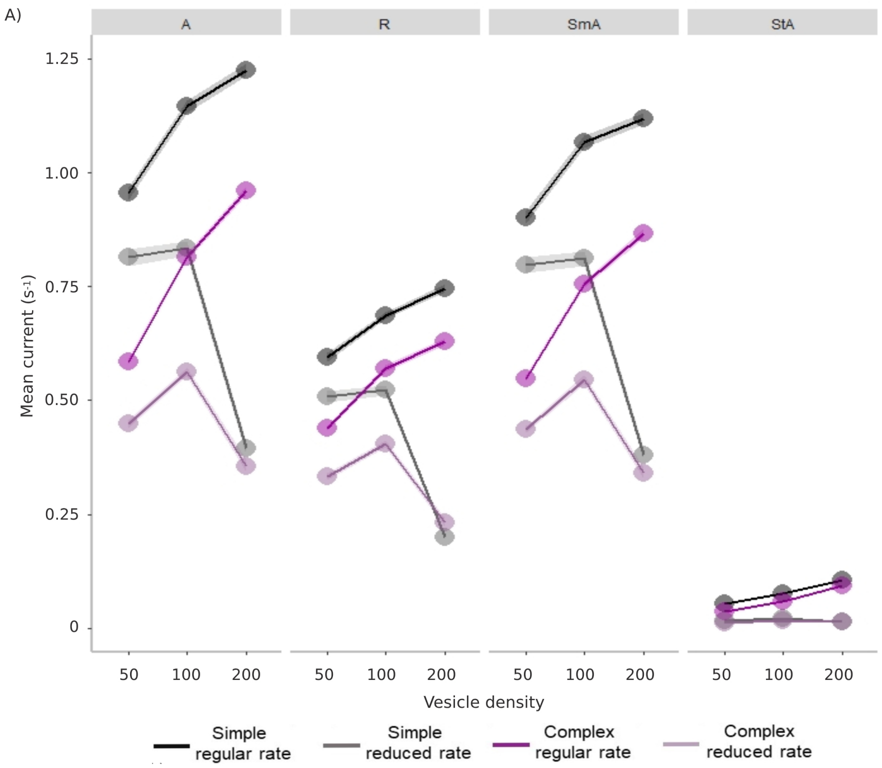


**Figure S4. Vesicle density and rates of switching between smooth and staggered motion states influence net anterograde vesicle transport**
The panel shows the smooth anterograde (SmA), step anterograde (StA), total anterograde (A), and retrograde (R) currents across different vesicle densities, at regular and reduced rates, for the simple and complex systems. Mean of 150 simulations ± 95% CI of the mean is plotted.

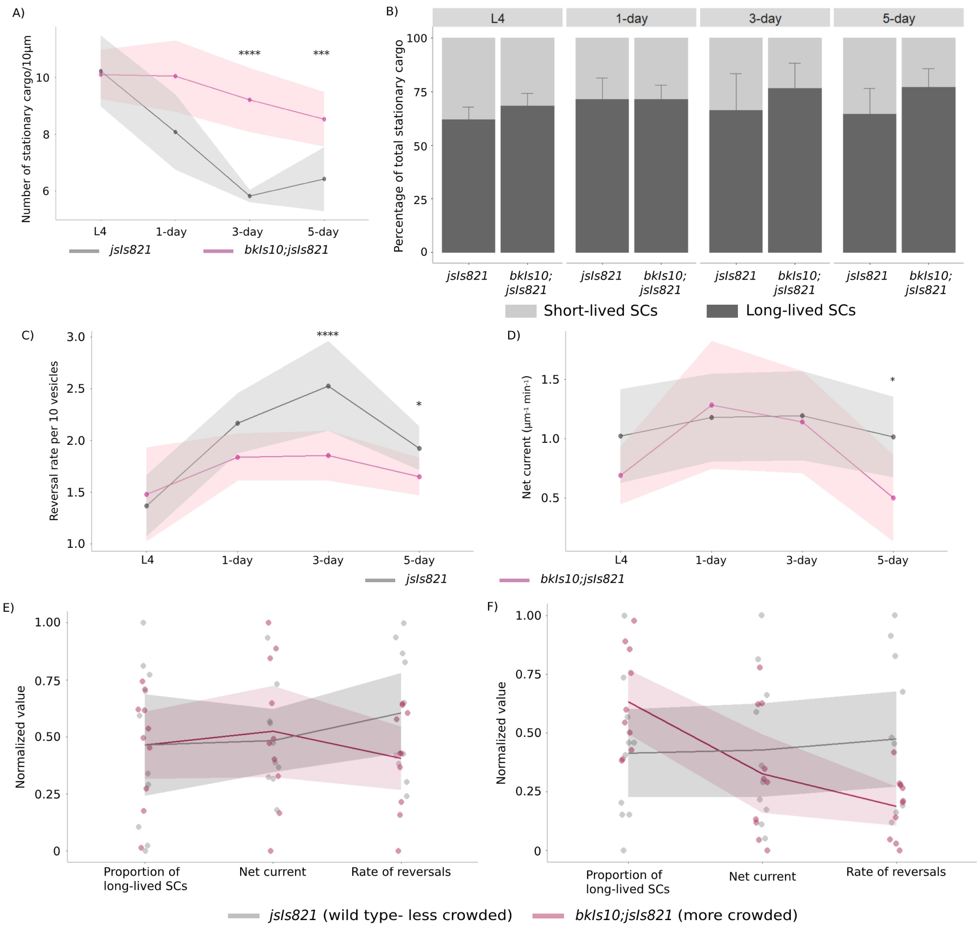
A)
Number of stationary cargo/10µm
****
***
L4
1-day
3-day
5-day
jsIs821
bkIs10;jsIs821
B)
L4
1-day
3-day
5-day
Percentage of total stationary cargo
jsIs821
bkIs10; jsIs821
Short-lived SCs
Long-lived SCs
C)
Reversal rate per 10 vesicles
D)
Net current (µm⁻¹ min⁻¹)
E)
Normalized value
Proportion of long-lived SCs
Net current
Rate of reversals
F)
jsIs821 (wild type- less crowded)
bkIs10;jsIs821 (more crowded)

**Figure S5**

**Figure S5. Age-dependent increase in crowding in TRNs of a tauopathy model of *C. elegans* is accompanied by a reduction in reversal rate and net anterograde vesicular transport**
(A) This graph compares the stationary cluster density (including both short-lived and long-lived SCs) between wild type (*jsIs821*) and a tauopathy model of *C. elegans* (*bkIs10; jsIs821*) across 4 stages (L4s, 1-day, 3-day, and 5-day adults). The solid circles and solid lines represent the mean values of stationary cluster density across stages, and the shaded bands represent the 95% CI around the mean. Mann-Whitney test conducted for test of statistical significance (***$p < 0.001$, ****$p < 0.0001$). N = 10 animals each for L4, 1-day and 5-day stage. L4: n (number of stationary vesicles) = 537 (*jsIs821*), 461 (*bkIs10;jsIs821*);1-day: n = 391 (*jsIs821*), 549 (*bkIs10; jsIs821*); 5-day animals: n = 277 (*jsIs821*), 370 (*bkIs10;jsIs821*). N = 16 animals for *jsIs821* 3-day stage, n = 426. N = 11 animals for *bkIs10; jsIs821* 3-day stage, n = 486.
(B) The graph compares the percentage distribution of short-lived and long-lived stationary clusters across stages between *jsIs821* and *bkIs10; jsIs821.* Mean ± SD plotted. N = 10 animals each for L4, 1-day and 5-day stage. L4: [nSL (number of short-lived stationary vesicles), nLL (number of long-lived stationary vesicles)] = [208, 329 (jsIs821)], [147, 314 (*bkIs10; jsIs821*)]; 1-day: [nSL, nLL] = [107, 284 (*jsIs821*)], [155, 394 (*bkIs10; jsIs821*)]; 5-day: [nSL, nLL] = [102, 175 (*jsIs821*)], [87, 283 (*bkIs10; jsIs821*)]. N = 16 animals for *jsIs821* 3-day, [nSL, nLL] = [143, 283]. N = 11 animals for *bkIs10; jsIs821* 3-day, [nSL, nLL] = [110, 376].
(C) This graph compares the reversal rate per 10 vesicles between *jsIs821* and *bkIs10; jsIs821* across stages. The solid circles and solid lines represent the mean values of reversal across stages, and the shaded bands represent the 95% CI around the mean. Mann-Whitney test conducted for test of statistical significance (****$p < 0.0001$, *$p < 0.05$). N = 10 animals each for L4, 1-day and 5-day stage. L4: n (number of reversals) = 644 (*jsIs821*), 387 (*bkIs10; jsIs821*); 1-day: n = 729 (*jsIs821*), 698 (*bkIs10; jsIs821*); 5-day: n = 582 (*jsIs821*), 324 (*bkIs10; jsIs821*). N = 16 *jsIs821* 3-day stage, n = 1237. N = 11 for *bkIs10; jsI821* 3-day stage, n = 557.
(D) This graph compares the net anterograde current between *jsIs821* and *bkIs10; jsIs821* across stages. The solid circles and solid lines represent the mean values of reversal across stages, and the shaded bands represent the 95% CI around the mean. Mann-Whitney test conducted for test of statistical significance (*$p < 0.05$). N = 10 animals each for L4, 1-day and 5-day stage. L4: n (number of moving vesicles) = 4794 (*jsIs821*), 2716 (*bkIs10; jsIs821*); 1-day: n = 3448 (*jsIs821*), 3734 (*bkIs10; jsIs821*); 5-day: n = 3064 (*jsIs821*), 1958 (*bkIs10; jsIs821*). N = 16 for *jsIs821* 3-day stage, n = 4813. N = 11for *bkIs10; jsI821* 3-day stage, n = 3059.
(E-F) These panels depict parallel plots that compares normalized values of the proportion of long-lived SCs, net current, and reversal rate between *jsIs821* and *bkIs10*; *jsIs821* in 1-day (E) and 3-day (F) adult *C. elegans* hermaphrodites. The number of animals and vesicles used in this analysis are the same as the ones described for panels (B-D).
