## Supplementary Information for "Cargo crowding, stationary clusters and dynamical reservoirs in axonal transport"

### The Monte Carlo Simulation Model

Our model for vesicle motion is framed directly in terms of vesicles and their dynamics, as inferred from experimental data. We model the axon as a set of “compartments”, each containing a fixed number of “tracks”, chosen to be 10 here. We work with a single compartment in this paper, representing the sub-region imaged in the experiments. Each model microtubule track is a one-dimensional line, discretized into independent, equally spaced lattice sites. Each lattice site is associated with at most a single vesicle. The discretization represents the basic 8 nm periodicity of the dimeric tubulin unit of the microtubule. We measure length in these units.

These tracks are aligned, as appropriate to the axon and arranged in a concentric ring, simulating those microtubules which lie towards the outside of the dense bundle within the touch neuron. The centres of vesicles are associated with a single site, excluding other vesicles from occupying the same site as well as adjacent ones. Since cargo dimensions are typically larger than the motor dimension, vesicles physically exclude each other over a larger scale. To model this, we assume that if the centre of a vesicle is associated with a specific site, it blocks two sites in front and two sites at the back from being occupied [1], [2], [3]. (If a vesicle is centred at site 4, the centre of another vesicle can only occupy sites 0 or 8 and no site in-between.) This corresponds to a vesicle diameter of about 3 X 8nm =

24nm, towards the lower end for estimates of vesicle size (approximately 16-48nm).

Visual analysis of kymographs from *in vivo* experiments imaging vesicle motion identifies three distinct motion states. Two of these states, smooth anterograde (SmA) and step anterograde (StA), refer to anterograde motion. SmA trajectories are relatively smooth whereas StA vesicles exhibit stop-and-go motion. Experiments in wildtype neurons show one generic state of retrograde motion whose properties qualitatively resemble vesicles with smooth anterograde motion (SmA). Vesicles exhibiting such smooth retrograde motion are labeled R. We prescribe hopping rates for these vesicles consistent with the properties of these states (Table S2).

Vesicle motion can be blocked by the presence of another vesicle on the same track. If a vesicle is blocked from motion, it can change its position or state. Its position can be changed through track-changing (equivalently, side-stepping - we use these terms interchangeably) events. Alternatively it can maintain its position but change its state.

In the "simple system", vesicles move on complete tracks and are only blocked by the presence of other vesicles in front of them. A more complex version of our model, the "complex system", incorporates further barriers to vesicle motion, over and above the blocking that arises from the presence of other vesicles. First, vesicle motion can be obstructed by the presence of microtubule ends. The number of such ends is that they occur at a frequency of about 1 per micron. We assume that when vesicles come to the end of a microtubule track on which they are traveling, they "sidestep" or "change state" with a small probability, but cannot advance on the same track. Second, a vesicle can be obstructed by larger physical obstructions to vesicle motion, such as mitochondria. Such mitochondria move relatively slowly on the scales of vesicle motion in a bi-directional fashion, but can be considered to be effectively stationary across the time-scales for vesicle motion. We assume that the presence of a mitochondrion results in a correlated block, in which sites with the same index on several adjacent microtubules (we assume 3 microtubules are collectively blocked) are prevented from being occupied. The rules by which vesicles move across these obstacles also involve "sidestepping" or "changing state" with a small probability, but the fact that several sites are blocked together implies that vesicle motion across these larger blocks is more strongly inhibited.

Our simulations must accommodate events which occur over a broad range of times scales, from the fast anterograde or retrograde motion of single vesicles to the relatively infrequent events in which vesicles change track or change state. We use a standard continuous time Gillespie algorithm [4] in which every allowed update (event) (a vesicle hopping, side-stepping or state-changing), occurs at a prescribed rate (see Table S2) $\gamma_i(\mathrm{t})$. The probability of the event occurring in a time interval dt about t i.e. [t; t + dt) is $\gamma_i(\mathrm{t})\mathrm{dt}$. Given these rates, the Gillespie method consists of calculating two stochastically generated quantities: the time at which the next event occurs and the specific update event. We measure all rates in units of $s^{-1}$, ensuring that our rates can be easily related to physically meaningful quantities [4].

We assume periodic boundary conditions, since these yield a steady-state current through the system without the need for assuming specific input and removal rates at the ends. Generalizations of our model to the case where we can allow for the entry and removal of vesicles at the two ends of the simulated compartment, maintaining the same density on average, yield similar results.

### Units and scales

At saturation ATP concentration, the mean velocity of smooth anterograde motion is of the order of 1 micron per second. A vesicle thus moves a distance of 8 nm in a time $\approx 10^{-2}s$, if unblocked. We will take this to set the basic hopping rate in our system: $100s^{-1}$. Thus, in a time of $10^4 \times 10^{-2}s = 10^2s$, or around a minute and a half, an isolated vesicle will be expected to have moved around 8000 sites or approximately $8 \times 10^{-9}m \times 10^4 = 80\mu m$ if it moved smoothly and continuously with fixed velocity. We assume, for simplicity, that the hopping rates for smooth (both retrograde and anterograde) motion are the same. For step motion, i.e. the StA state, we choose this hopping rate to be $40s^{-1}$. This ensures that, compared to the smooth states, the staggered state pauses more often and for longer.

We choose rates for side-stepping and state changing, to be much smaller, consistent with the presence of substantial stationary clusters of vesicles in the experiments - had these been substantially larger, we would see virtually no long-live clusters. *In vitro* measurements of dynein and kinesin coated beads suggests that cytoplasmic dynein exhibits more substantial off-axis movements (5 times per micron) as opposed to kinesins (0.9 times per micron) [5]. Given that a processive motor can walk upto a micron or more without detachment, this suggests that, provided we interpret off-axis movements as equivalent to changing protofilament tracks, side-stepping rates can reasonably be expected to be around a hundred to a thousand times smaller than rates for motor advance i.e. two vesicles encountering each other on the same track should relax relatively slowly through changes of track.We choose these to be $0.02s^{-1}$ for all vesicle states; for the sake of consistency between the experimental and simulation bleach recoveries. Finally, for the interconversion rates between vesicles that are stalled, in steady state, decouples from the other rates, we choose these to be of order $0.1 - 1s^{-1}$ . But these numbers depend on the specific vesicle state and differ between SmA, StA and R, so must be chosen depending on the specific transition.

We make several approximations to avoid the proliferation of rates that cannot be measured directly, although our simulations can allow all relevant rates to vary, in principle. If we make the reasonable assumption that SmA and R correspond to an excess of anterograde retrograde motors respectively, with the step state corresponding to a roughly equal distribution of + and -end directed motors, direct transitions between the smooth anterograde and smooth retrograde states can be expected to be unlikely, since this would require the coordinated attachment/detachment of a number of motors simultaneously. Thus we only allow transitions between the SmA and StA and between R and StA separately, but not the direct SmA to R transition. Given the relatively smaller number of StA vesicles, we argue that the rates for transitions out of the StA state should be larger than the rates for transitions into this state. Solving the master equation for the relative fractions of SmA, StA and R states yield an initial choice of rates, which we then further optimise against the experimental ratio of SmA:StA:R (50:10:40). Our final rates are listed in the Table S2.

Vesicles do not enter or leave our system overall, given periodic boundary conditions, and we thus work with a fixed number of vesicles (50-200) across 1000 sites X 10 microtubules i.e. 10000 sites in all. Our model thus simulates an 8micron linear section of the axon, comparable to the region imagined. Our vesicle densities are then 6.3, 12.5, 25 vesicles per micron. Experimental numbers are believed to be of comparable order. We simulate a maximum of 10 full-length microtubules, although we can break these into smaller sections to simulate the effects of microtubule ends on transport. For simulating photobleaching and axotomy are simulated on a 64 $\mu m$ system with an initial vesicle density of 400, 800 and 1000 vesicles. The vesicle numbers correspond to densities of 6.3, 12.5, 15.6 vesicles per micron. An 8 $\mu m$ region was bleached in the photobleaching simulation and the ablated region spanned 0.8 $\mu m$ in the Axotomy simulation.

To simulate the effects of data sampling through discrete snapshots (frames), we represent data in our simulation kymographs in terms of snap- shots separated by 0.2s. A 5 fps camera takes a snapshot at every $2 \times 10^{-1} s$,yielding an equivalent sampling time. Thus, when we compare kymographs generated through our methodology with experimental kymographs, we accumulate static images separated by this time period.

### What is measured

We simulate long enough that the system reaches steady state, allowing for a large number of updates (typically $10^7$ or more) to elapse before the system can be assumed to reach steady state. We test for steady state in several ways. The current must be constant throughout the system and we compare the averaged flux at 3 equally spaced locations along the full length of the axon to ensure this. We also compute the averaged number of smooth, step and retrograde movers, checking that they have converged to time-independent values.

#### Current:

The current (equivalently the flux) is obtained by computing the time-averaged currents of different states of vesicles in the following way: Choosing a link connecting two neighbouring sites, we compute the number of anterograde vesicles of both states (smooth and step) moving across it, as well as the number of retrograde vesicles, over an interval of time. Subtracting the numbers of retrograde vesicles from anterograde vesicles, and dividing by the elapsed time, gives us the current, which can then be computed over a large number of configurations to generate averages in steady state.

#### Relative Proportion of Vesicles:

Experimental observations indicate that 51 to 58 percent of vesicular traffic is comprised of smooth anterograde vesicles, while step anterograde comprises 4 to 11 percent of the traffic and retrograde movers make up the remaining 37 to 43 percent. In our model, the rates at which vesicles change states are chosen so as to produce a comparable population of vesicles overall. We monitor the relative proportion of SmA:StA:R along the axon, confirming that the StA state is generated predominantly at clusters.

#### Cluster Lifetimes:

We track each individual vesicle, so we can easily measure the statistics, averaged over all vesicles in steady state, of stationary vesicles in clusters. In experiments, individual vesicles cannot be identified once they enter stationary clusters, the number of vesicles trapped in any given cluster cannot be accessed directly and individual microtubules cannot be separately resolved. This implies that we must average over all microtubules at a fixed location, to compare the predictions and kymographs generated by the model to the experimental data. Our definition of a cluster must also incorporate the fact that, since the camera frame-rate is 5 fps, we should not resolve motion at faster scales in the simulation if we are to compare our results to experimental data. We take snapshots of the axon at time intervals which correspond to the camera frame rate, as discussed above. A site is considered to be occupied, contributing to the fluorescence signal, if the corresponding site in any of the microtubules is occupied. We compute the duration of the clusters and the frequency of their occurrence.

#### Number of long-lived stationary clusters per 8 microns:

We count each cluster and its duration through the entire length of the simulation, resolving both relatively short-lived clusters arising from the collisions of pairs of oppositely directly vesicles as well as longer-lived ones arising from the piling up of vesicles at simulated microtubule ends and blocks associated with mitochondria.

#### Kymographs

To construct kymographs from vesicle configurations in our data, we must reexamine how such kymographs are extracted from the experimental movies. First, moving vesicles show up as fainter lines on the kymographs. Second, both the camera and the subsequent analysis software will tag a labeled vesicle only if its intensity exceeds a threshold. Third, the camera images vesicles at a 5 fps rate. Thus we should connect vesicle trajectories over intervals of 0.2s, as earlier calculated, in order to accurately relate our data to the experiments. Given this, we choose the following algorithm for generating our kymographs. We plot all vesicles. In our kymographs, there are 2 parameters for the trajectories, the greyscale value and the thickness. Of all the vesicles, we identify those who are part of long-lived clusters, representing them by thicker (darker) lines in the kymograph.

The grey-scale is assigned in proportion to the time spent by such vesicles in stationary clusters. To set these (thickness and greyscale) values, we compute the maximum time each vesicle spends at a site. If this is 0.6s or 0.8s, the thickness is set to 3, 4 respectively. For all other values of maximum time (0.2s, 0.4s, 1s, 1.2s...), the thickness is 1. For the greyscale value, if the maximum time is $< 1s$, a lighter shade of grey is used (RGB value [0.3, 0.3, 0.3]), while for values of maximum time $> 1s$, a darker shade of grey is used (RGB value [0.1, 0.1, 0.1]).

| Strain name | Description | Experiments |
|---|---|---|
| *jsIs821* ( [6, 7]) | *mec-7p::GFP::RAB-3* | Live imaging for obtaining currents, stationary cluster lifetimes, reversal rates, and area occupied by stationary clusters<br>Fluorescence Recovery after Photobleaching |
| *jsIs37* ( [8]) | *mec-7p::SNB-1::GFP* | Axotomy |
| *bkIs10;jsIs821* ( [9], this study) | *bkIs10[aex-3p::Tau-337M myo-2p::GFP] jsIs821[mec-7p::GFP::RAB-3]* | Live imaging for obtaining currents, stationary cluster lifetimes, and reversal rates |

**Table S1. List of strains used in this study**

| **Parameter** | **Value ($s^{-1}$)** |
|---|---|
| Hopping rate for Smooth Anterograde vesicles (SmA) | 100 |
| Hopping rate for Staggered Anterograde vesicles (StA) | 40 |
| Hopping rate for Smooth Retrograde vesicles (SmR) | 100 |
| Sidestepping rate for Smooth Anterograde vesicles (SmA) | 0.02 |
| Sidestepping rate for Staggered Anterograde vesicles (StA) | 0.02 |
| Sidestepping rate for Smooth Retrograde vesicles (SmR) | 0.02 |
| Conversion rate for SmA $\rightarrow$ StA | 0.08 |
| Conversion rate for SmA $\rightarrow$ SmR | 0 |
| Conversion rate for StA $\rightarrow$ SmA | 0.8 |
| Conversion rate for StA $\rightarrow$ SmR | 0.64 |
| Conversion rate for SmR $\rightarrow$ StA | 0.08 |
| Conversion rate for SmR $\rightarrow$ SmA | 0 |

**Table S2. Choice of Simulation Rates**
The choice of rates used in our kinetic Monte Carlo simulations, described in more detail in Supplementary Information. Each rate is provided in natural units of $s^{-1}$. The ratio of rates provided, indicates for example, that we take the rates of side-stepping of an anterograde vesicle to be about a 1000 times smaller than the rate at which it moves forward if unblocked.

**Movie S1 Influence of microtubule track switching events on cargo transport** The movie shows the evolution of the simulation system in which initially both side-stepping and reversals of vesicle cargo are disallowed. This leads to a jammed configuration with zero net current at around 15 seconds as the vesicles cannot circumvent obstacles by side-stepping on to a neighbouring microtubule track or reverse their direction of motion. This point on, vesicles are allowed to side-step on to a neighbouring microtubule track but not reverse their direction of motion on encountering an obstacle. Transport resumes but the system eventually reaches a permanent jammed state.

**Movie S2 Influence of vesicle reversal events on cargo transport** The movie shows the evolution of the simulation system in which initially both side-stepping and reversals of vesicle cargo are disallowed. This leads to a jammed configuration with zero net current at around 19 seconds as the vesicles cannot circumvent obstacles by side-stepping on to a neighbouring microtubule track or reverse their direction of motion. This point on, vesicles are allowed to reverse their direction of motion but not side-step on to a neighbouring microtubule track on encountering an obstacle. Transport is restored in the system.
